# Hedgehog signaling is required for endomesodermal patterning and germ cell development in *Nematostella vectensis*

**DOI:** 10.1101/2020.01.15.907238

**Authors:** Cheng-Yi Chen, Sean A. McKinney, Lacey R. Ellington, Matthew C. Gibson

## Abstract

Two distinct mechanisms for primordial germ cell (PGC) specification are observed within Bilatera: early determination by maternal factors or late induction by zygotic cues. Here we investigate the molecular basis for PGC specification in *Nematostella*, a representative pre-bilaterian animal where PGCs arise as paired endomesodermal cell clusters during early development. We first present evidence that the putative PGCs delaminate from the endomesoderm upon feeding, migrate into the gonad primordia, and mature into germ cells. We then show that the PGC clusters arise at the interface between *hedgehog1* and *patched* domains in the developing mesenteries and use gene knockdown, knockout and inhibitor experiments to demonstrate that Hh signaling is required for both PGC specification and general endomesodermal patterning. These results provide evidence that the *Nematostella* germline is specified by inductive signals rather than maternal factors, and support the existence of zygotically-induced PGCs in the eumetazoan common ancestor.

## Introduction

During development, animal embryos typically set aside a group of primordial germ cells (PGCs) that later mature into germline stem cells (GSCs) and in turn give rise to gametes during adulthood (Nieuwkoop and Sutasurya, 1979, 1981; Wylie, 1999; Juliano, Swartz and Wessel, 2010). The process of PGC specification both underpins the sexual reproduction cycle and involves transitions of pluripotency, making the mechanisms that distinguish germ cells from soma of critical importance in developmental and stem cell biology (Solana, 2013; Irie, Tang and Azim Surani, 2014; Magnúsdóttir and Surani, 2014). Comparative studies on germ cell development have defined two common mechanisms of PGC specification among diverse animals: preformation and epigenesis (Nieuwkoop and Sutasurya, 1979, 1981; Extavour and Akam, 2003). During PGC specification by preformation (e.g. *Drosophila*, *C. elegans* and *Danio rerio*), cytoplasmic determinants referred to as the germ plasm are maternally deposited into embryos and then segregated into specific blastomeres through cell division (Strome and Wood, 1982; Williamson and Lehmann, 1996; Yoon, Kawakami and Hopkins, 1997). In contrast, there are neither maternal germline determinants nor pre-determined PGC fates in specific blastomeres in epigenic PGC specification. For example, BMP signaling is required for PGC specification from precursor cells in mouse, axolotl, and cricket embryos (Lawson *et al*., 1999; Chatfield *et al*., 2014; Nakamura and Extavour, 2016). The epigenesis mode of PGC specification is more prevalent across the animal kingdom, and therefore hypothesized to reflect mechanisms present in the cnidarian-bilaterian common ancestor (Extavour and Akam, 2003). However, to date, no mechanistic studies of PGC specification in early branching animals have functionally tested this hypothesis.

Cnidarians (jellyfish, sea anemones and corals) are the closest sister group to bilaterians and occupy an ideal phylogenetic position for investigating likely developmental traits of the eumetazoan common ancestor (Technau and Steele, 2011; Russell *et al*., 2017). Among cnidarians, the sea anemone *Nematostella vectensis* maintains distinct adult gonad tissue and features PGC specification dynamics hypothesized to partially reflect ancient epigenesis based on expression patterns of conserved germline genes (Extavour *et al*., 2005). Additionally, a well-annotated genome (Putnam *et al*., 2007), defined developmental stages (Fritzenwanker *et al*., 2007) and diverse genetic tools (Ikmi *et al*., 2014; Renfer and Technau, 2017; He *et al*., 2018; Karabulut *et al*., 2019) make *Nematostella* a genetically tractable model to elucidate developmental mechanisms controlling PGC specification.

In this study, we explore mechanisms of PGC development in *Nematostella* and test whether the putative PGC clusters are specified by maternal or zygotic control. We first follow the development of putative PGCs and provide evidence supporting their germ cell fate in adults. We then leverage shRNA knockdown and CRISPR/Cas9 mutagenesis to interrogate the developmental requirements for the Hedgehog signaling pathway in PGC specification. From these results, we conclude that Hh signaling is either directly or indirectly required for PGC specification in *Nematostella*. As Hh signaling is only activated zygotically, these data indicate an epigenic mechanism for *Nematostella* PGC specification and support the inference that the eumetazoan common ancestor specified PGCs via epigenesis.

## Results

### Evidence that PGCs form in primary polyps and migrate to gonad rudiments

The localized expression of the conserved germline genes *vasa*, *nanos*, and *piwi* suggest that *Nematostella* PGCs arise within two cell clusters of the pharyngeal endomesoderm of primary polyps (Fig. 1, Fig. S1; Extavour *et al*., 2005; Praher *et al*., 2017). To follow the development of putative PGCs at higher spatio-temporal resolution, we generated a polyclonal antibody against *Nematostella* Vasa2 (Vas2) and used immunohistochemistry and fluorescent *in situ* hybridization to confirm that Vas2 was co-expressed with *piwi1* and *piwi2* in the putative PGC clusters (Fig. S1 E-L). Further supporting their germline identity, we also found that *tudor* was enriched in putative PGC clusters (Fig. S1 M-P). To gain more detailed spatial information, we reconstructed confocal z stack images in 3D and found that Vas2+ epithelial cell clusters were localized at endomesodermal junctions where the pharyngeal endomesoderm connects to the primary mesenteries (Fig. 2B-B’; Sup. Mov.1).

**Fig. 1.**
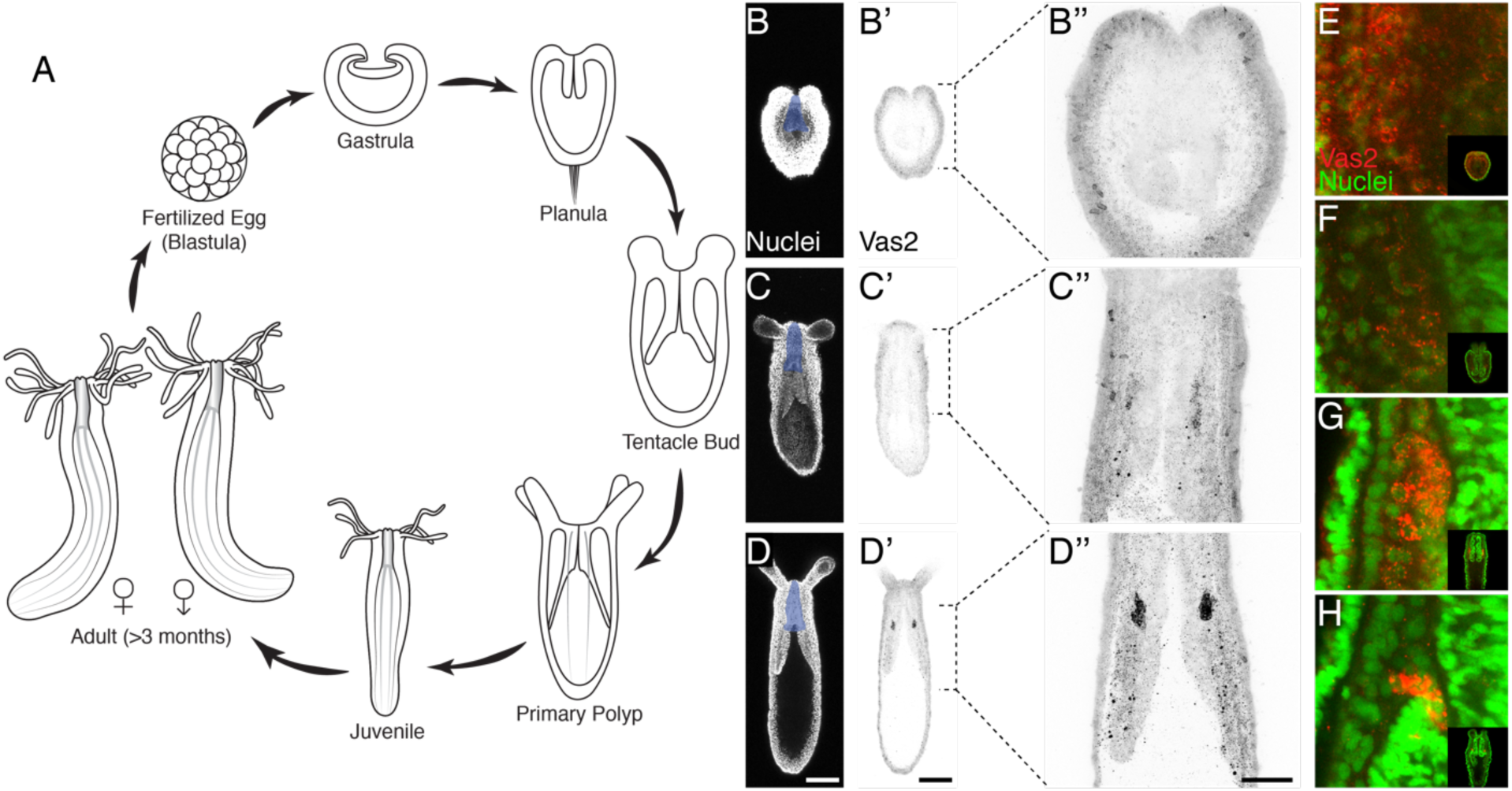
Putative *Nematostella* PGCs clusters are specified during metamorphosis. (**A**) Diagram depicting the *Nematostella* life cycle. Nuclei staining with Hoechst (**B-D**) and Vas2 immunostaining (**B’-D’** and **B’’-D’’**) during tentacle bud to primary polyp metamorphosis. Vas2 expression is gradually enriched at two cell clusters next to the pharynx (shaded *blue*), showing PGC specification in certain endomesodermal cells. (**E-H**) The maternally-deposited Vas2 protein (*red*) forms granules around the nuclei of endomesodermal cells (*green*), which diminish as the PGC clusters form. Scale bar = 100 µm in **D-D’**; 50 µm in **D’’**; **B-D’** are at the same scale; **B’’-D’’** are at the same scale; **E-H** are at the same scale.

**Fig. 2.**
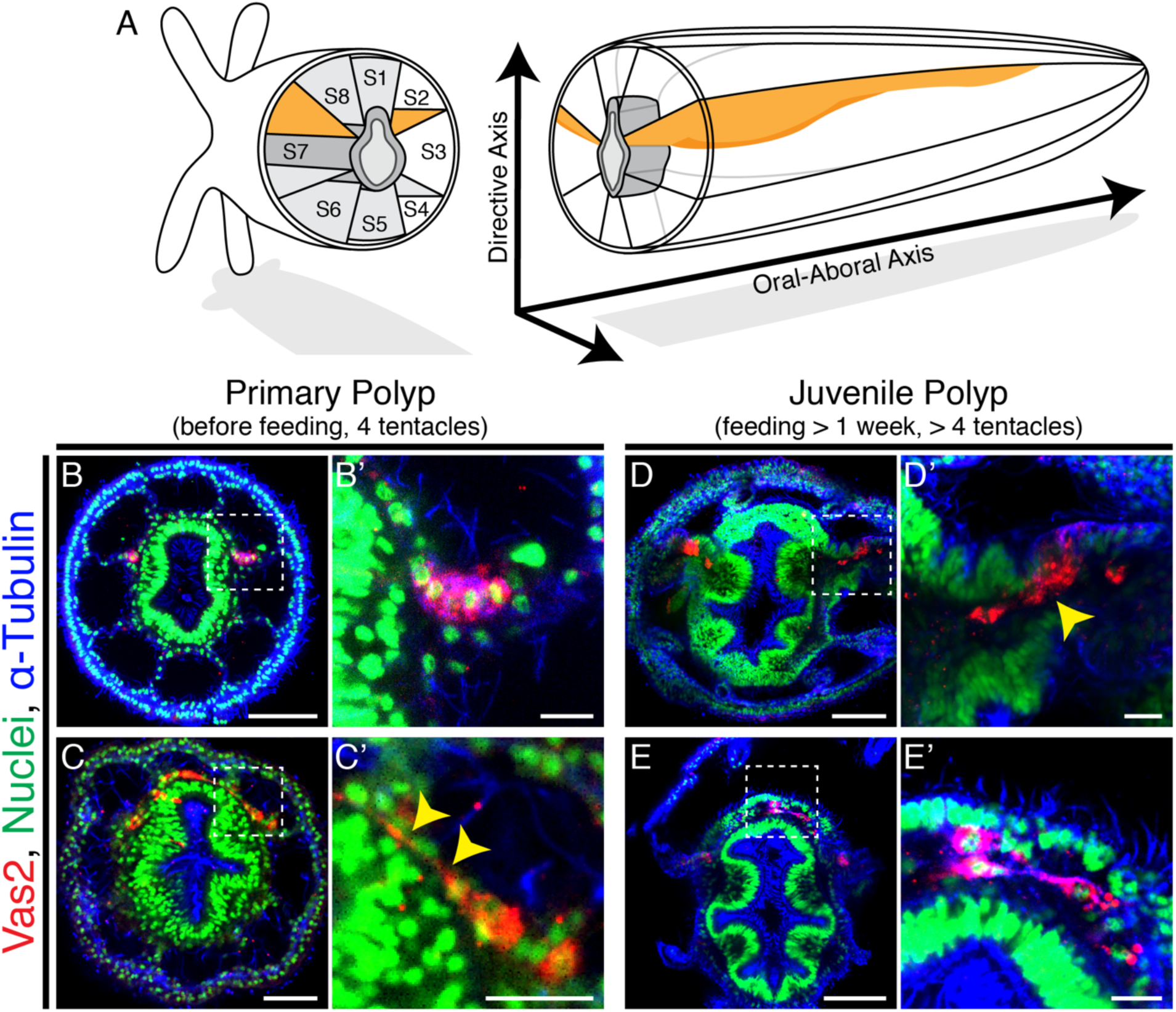
Putative *Nematostella* PGCs delaminate through epithelial-mesenchymal transition (EMT) and appear to migrate to non-primary mesenteries. (**A**) Schematic diagram of *Nematostella* polyp anatomy depicts the pharynx and mesentery arrangements at the pharyngeal level. The eight mesenteries (two primary mesenteries in *orange* and six non-primary mesenteries in *light gray*) harbor gonad epithelium, muscle and digestive tissue. The internal structures of *Nematostella* are arrayed around the pharynx (*dark gray*). The *directive* and *Oral-Aboral* axes are indicated, segment nomenclatures follow He et al. (2018). (**B-B’**) Paired clusters of putative PGCs labeled by Vas2 immunofluorescence (*red*) initially exhibit epithelial charateristics. (**C-C’**) Putative PGCs from >10 dpf primary polyps appear to stretch their cell bodies basally (*yellow arrowheads*). (**D-D’**) Following nutrient intake, putative PGCs delaminate into the mesoglea through an apparent EMT (*yellow arrowhead*). (**E-E’**) In the mesoglea, these Vas2+ cells exhibit fibroblast-like morphology and are detected between mesenteries at the level of the aboral pharynx. Scale bar = 10 µm in **B’**, **C’**, **D’**, **E’**; 20 µm in **C**; 50 µm in **B**, **D**, **E**.

Adult *Nematostella* harbor mature gonads in all eight internal mesentery structures (Fig. S2A-C E-E’; Williams, 1975; Frank and Bleakney, 1976). If the two Vas2+ epithelial cell clusters are the only precursors for adult germ cells, it follows that these cells would have to delaminate and migrate to populate the eight gonad rudiments. Alternatively, new PGCs could arise within each of the six non-primary mesenteries, perhaps at a later developmental stage. To distinguish between these possibilities, we examined the localization of Vas2-expressing putative PGCs in primary polyps and later juvenile stages. In the majority of primary polyps, putative PGCs initially appeared in two coherent clusters at 10 days-post-fertilization (dpf, Fig. 2B-B’, Fig. 3A-A’). In older primary polyps (>10 dpf), some PGC clusters cells appeared to stretch basally through the underlying cell-free mesoglea (Fig. 2C-C’). After feeding for more than a week, primary polyps start adding tentacles and enter the juvenile stage. Interestingly, upon feeding, Vas2-expressing putative PGCs appeared to delaminate from the epidermis into the underlying mesoglea (Fig. 2D-D’). Delaminated Vas2 positive cells displayed a fibroblast-like morphology with filopodial protrusions, similar to other migratory cell types (Fig. S3; Scarpa and Mayor, 2016). Consistent with migratory potential, Vas2+ cells also expressed *twist* (Fig. S1Q-T), a conserved regulator of mesoderm development and a marker of metastatic cancer cells (Yang *et al*., 2004; Kallergi *et al*., 2011).

**Fig. 3.**
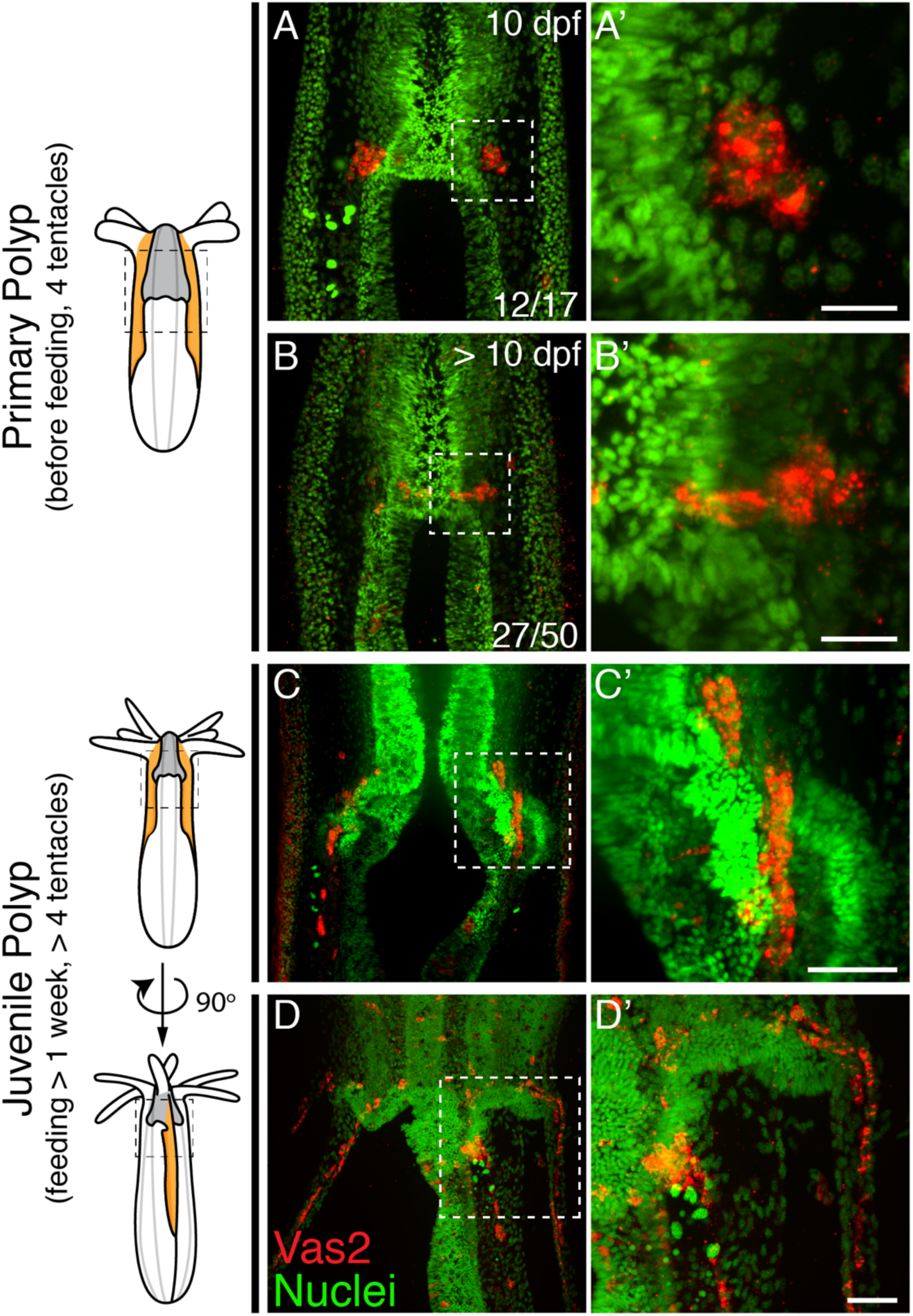
*Nematostella* PGCs migrate aborally to the gonad rudiments during juvenile stage. (**A-A’**) The majority of young primary polyps (≤10 dpf) exhibit two PGC clusters (Vas2+, *red*) in close proximity to the developing pharynx. (**B-B’**) In more mature primary polyps (>10 dpf), some PGCs elongate and localize between mesenteries. (**C-C’**) Following feeding, putative PGCs spread aborally into gonad rudiments. (**D-D’**) A juvenile polyp viewed 90 degrees from the orientation of **C**, showing aborally migrating PGCs in non-primary mesenteries. Scale bar = 10 µm in **A’**, **B’**; 20 µm in **C’**, **D’**.

We next followed the localization of putative PGCs through successive developmental timepoints. While specification of additional PGC clusters was not observed, we did find evidence for a process of radial cell migration between mesenteries at the level of the aboral end of the pharynx, where the mesoglea between ectoderm and endomesoderm increases in volume after the primary polyp stage (Fig. 2E-E’). The majority of 10 dpf primary polyps showed PGCs within clusters, while >10 dpf primary polyps showed some PGCs localized between the primary mesenteries and segment S1 (Fig. 2C, Fig. 3B-B’; He *et al*., 2018). The direction of this initial migration was away from the high BMP activity domain along the directive axis (Genikhovich *et al*., 2015), suggesting the potential existence of attractive/repulsive signals for migratory PGCs. Additionally, we found that putative PGCs migrated aborally from the clusters at the pharyngeal level to the mesenteries, where gonad rudiments will mature in adults (Fig. 3C-D’). Combining all observations, we hypothesize that in *Nematostella*, putative PGCs initially form as two endomesodermal cell clusters at the level of the aboral pharynx, delaminate into the cell-free mesoglea layer between the ectoderm and endomesoderm possibly via epithelial-mesenchymal transition (EMT), and then migrate to the gonad rudiments during juvenile development.

### Evidence that putative PGCs mature and give rise to germ cells in adult gonads

To assess the germline identity of putative PGCs, we next followed the development of Vas2+ cells from juvenile stages to young adults (>2-months old). In maturing polyps, the endomesodermal mesenteries are organized from proximal (external) to distal (internal) into parietal muscles, retractor muscles, gonads and septal filaments, with occasionally observed ciliated tracts between the gonads and septal filaments (Fig. 4A-D, Fig. S2H; Williams, 1979; Jahnel, Walzl and Technau, 2014). In juvenile polyps, Vas2+ cells were observed in the mesoglea between the septal filaments and the retractor muscles (Fig. 4C), an endomesodermal region that will later form adult gonad epidermis (Fig. S2H). We also occasionally observed putative PGCs between the ciliated tracts and the retractor muscles (Fig. 4D). After feeding for 8 weeks, most polyps reached the 12-tentacle stage and the mesenteries progressively matured, becoming wider and thicker. At this stage we observed Vas2+ putative PGCs in the maturing gonad region, along with Vas2+ immature oocytes or sperm cysts in females and males, respectively (Fig. 4E-F, Fig. S4). In adult mesenteries, similar putative PGCs were found in the vicinity of oocytes and sperm stem cells, as well as the aboral domains of the mesenteries (Fig. S2). Taken together, these observations suggest that the putative PGCs comprise a continuous Vas2-expressing lineage that proliferates and ultimately gives rise to mature germ cells for *ex vivo* sexual reproduction (Fig. S5, S6, Sup. Mov. 2). As proposed by previous work (Extavour *et al*., 2005), these data support the hypothesis that the germline gene-expressing cell clusters of primary polyps represent *bona fide* PGCs of *Nematostella*.

**Fig. 4.**
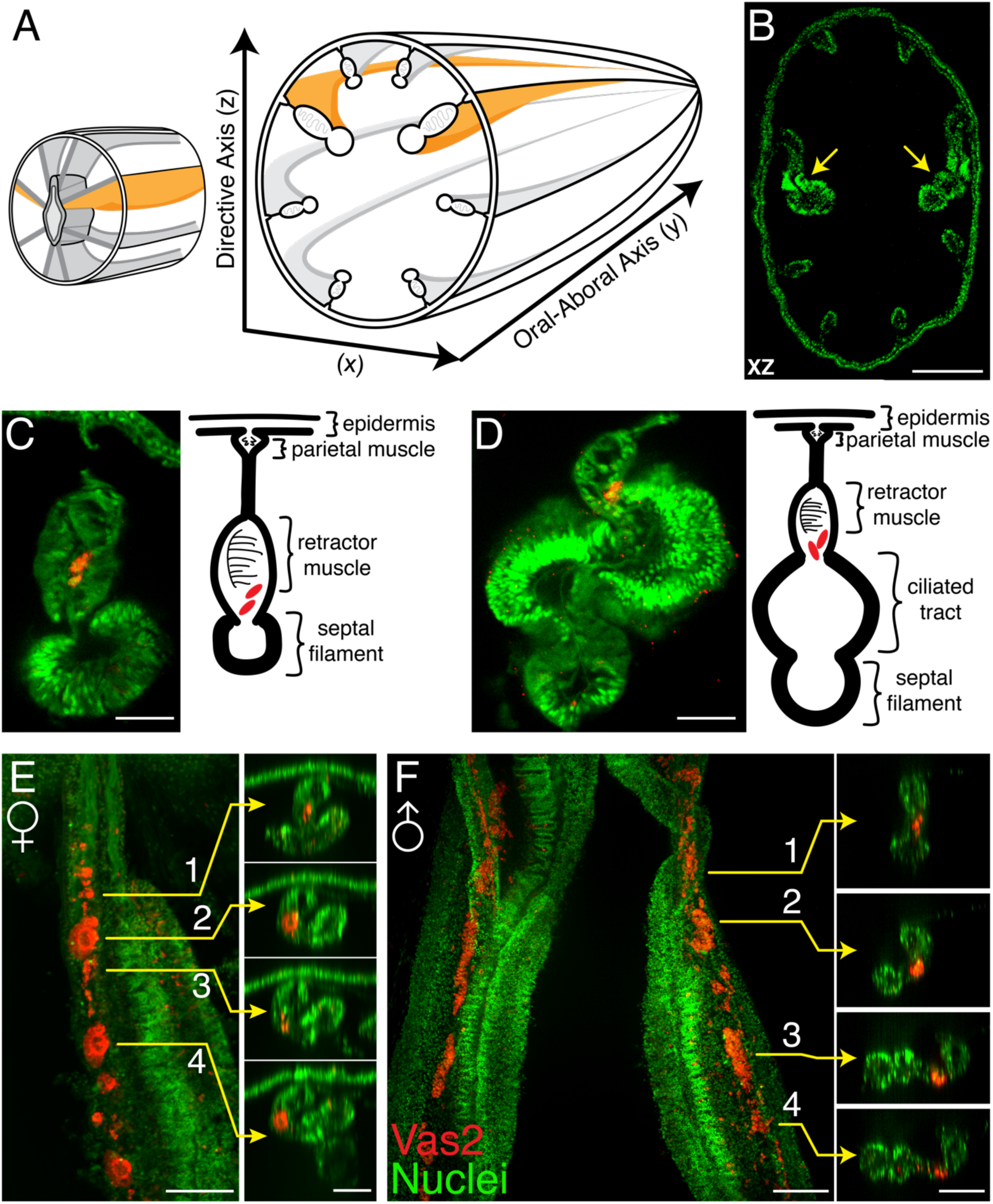
Vas2+ germ cells in juvenile gonad rudiments. (**A**) Schematic diagram of *Nematostella* polyp anatomy depicts the gametogenic mesenteries at the mid-body level. (**B**) A mid-body level cross section through a juvenile polyp, note the enlarged primary mesenteries (*yellow arrows*). Nuclei are counter stained by DAPI (*green*). (**C-D**) Representative images of maturing mesenteries with corresponding schematic diagrams. Putative PGCs are labeled by Vas2 immunofluorescence in *red*. (**E**) Whole-mount juvenile female mesentery shows Vas2-labeled putative PGCs and germ cells in close proximity (*red*) suggesting maturing oocytes originate from the continuous PGC lineage. (**F**) Whole-mount juvenile male mesentery shows Vas2-labeled putative PGC and GC populations, including the rudimentary sperm cysts. Insets 1-4 of **E** and **F** are xz plane views at the indicated levels. Scale bar = 100 µm in **B**; 20 µm in **C-F**.

### Evidence supporting a zygotic mechanism for primordial germ cell specification

We next investigated whether developing *Nematostella* form PGCs by a maternal preformation program or by a zygotically-driven epigenic process. In preformation, maternally-deposited germline determinants are segregated into specific PGC precursors during early cleavage (Nieuwkoop and Sutasurya, 1979, 1981; Extavour and Akam, 2003). In *Nematostella*, prior to the appearance of putative PGC clusters in developing polyps, we observed maternally-deposited perinuclear Vas2 granules that could hypothetically serve as germline determinants (Fig. 1E). These granules were previously identified with an independent antibody and proposed to regulate *Nematostella* piRNAs (Praher *et al*., 2017). However, perinuclear Vas2 granules were not restricted to a set of germline precursor cells and were distributed homogenously around oocyte germinal vesicles (Fig. S2B), in every cell of blastulae, and in most endomesodermal cells after gastrulation (Fig. 1E; Praher *et al*., 2017). In endomesodermal cells, Vas2+ granules gradually diminished when the putative PGC cluster cells activated Vas2 expression (Fig. 1E-H), suggesting germ cell fates were gradually specified from among endomesodermal precursor cells rather than being maternally pre-determined. Furthermore, germline gene transcripts (i.e. *vas1, vas2, nos2* and *pl10*) displayed a homogenous distribution in the endomesoderm of embryos and larvae before PGC specification (Extavour *et al*., 2005). Endomesodermal enrichment of “germline” genes before *Nematostella* PGC formation could be consistent with the proposed germline multipotency program (GMP), where the expression of conserved germline genes underlies multipotency of progenitor cells (Juliano, Swartz and Wessel, 2010). In line with GMP hypothesis, we hypothesize that *Nematostella* PGCs are zygotically specified from a pool of multipotent endomesodermal precursors.

In primary polyps, putative PGC clusters initially form in the two primary mesenteries, which are distinguished by the presence of aborally-extended regions of pharyngeal ectoderm known as septal filaments (Sup. Mov. 1, Fig. S7A-A’). While the mechanism of primary mesentery specification is unknown, this process likely lies downstream of Hox-dependent endomesodermal segmentation in developing larvae. Interestingly, segmentation of the presumptive primary mesenteries is disrupted in both *Anthox6a* mutants and *Gbx* shRNA-KD polyps (He et al., 2018). In both loss of function conditions, we observed aberrant attachment of the septal filaments and the associated induction of PGC clusters in non-primary septa (Fig. S7B-D’). This suggests that the precise location of the putative PGC clusters can be subject to regulation, and hints at the existence of zygotic PGC-inducing signals from the pharyngeal ectoderm.

### PGC specification is dependent on zygotic Hedgehog signaling activity

Previous gene expression studies have suggested that the Hh signaling pathway may be involved in patterning the endomesoderm and potentially the formation of germ cells (Matus *et al*., 2008). Using double fluorescent *in situ* hybridization to detect the expression of *Nematostella hedgehog1* (*hh1*) and its receptor *patched* (*ptc*) in late planula larvae, we found that both ligand and receptor were expressed in reciprocal domains of ectoderm and endomesoderm associated with the pharynx (Fig. 5A). Later, the PGC clusters appeared within the endomesodermal *ptc* expression domain, adjacent to where *hh1* is expressed in the pharyngeal ectoderm (Fig. 5B-D). Because PGCs formed in association with the juxtaposed *hh1* and *ptc* expression domains, we hypothesized that Hh signaling may direct neighboring endomesodermal cells to assume PGC identity (Fig. 5E).

**Fig. 5.**
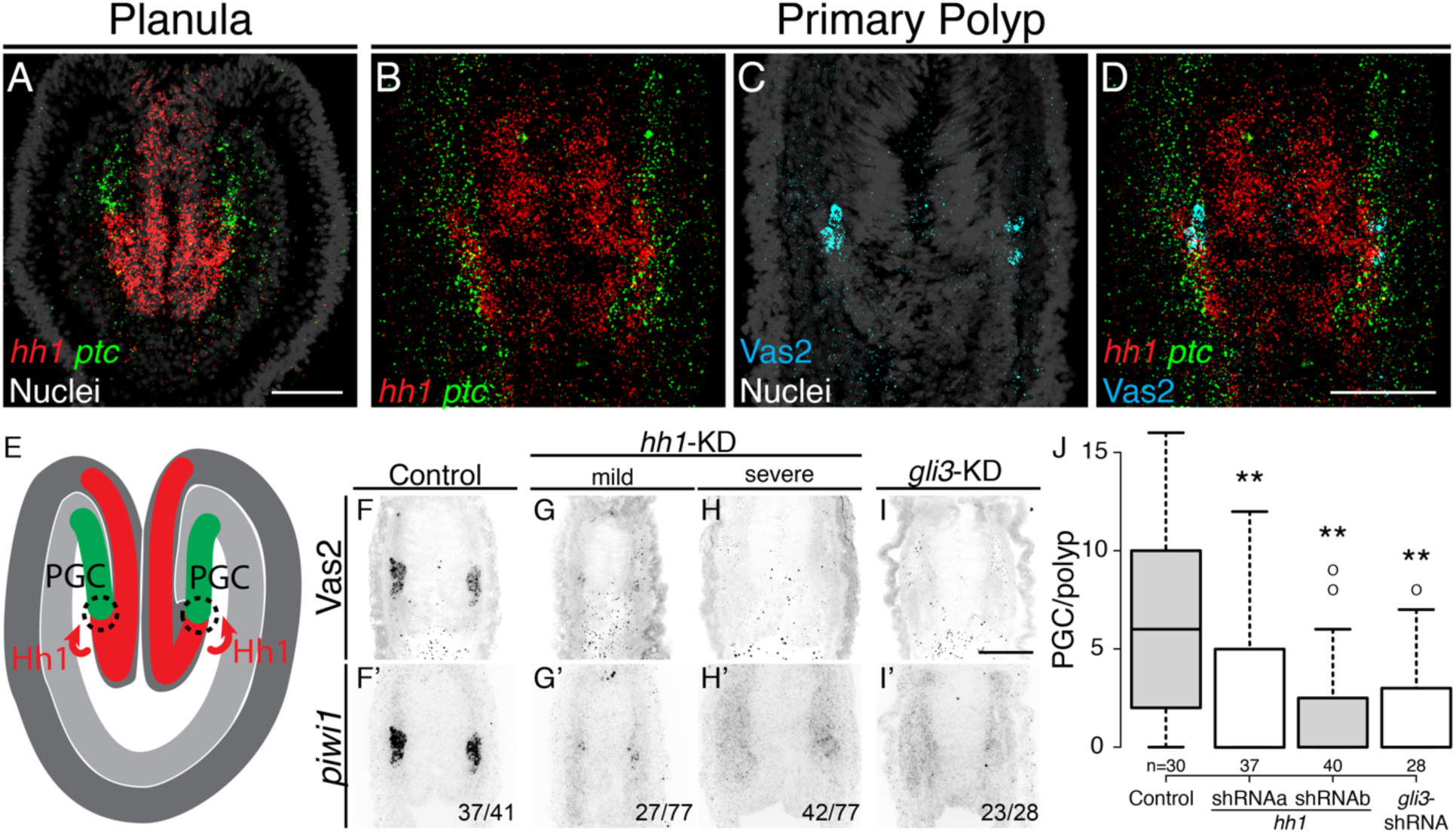
Hh signaling is required for *Nematostella* PGC formation. (**A**) Prior to PGC specification in planula larvae, *hh1* (*red*) and *ptc* (*green*) are expressed in pharyngeal ectoderm and endomesoderm, respectively. (**B-D**) In primary polyps, PGC clusters (*cyan*) are specified within the *ptc* expression domain, neighboring to the *hh1* domain. (**E**) Diagram depicting our working hypothesis about how Hh regulates PGC formation. (**F-I’**) PGC formation—indicated by Vas2 immunostaining and *piwi1* fluorescent *in situ* hybridization—is impaired by *hh1* or *gli3* shRNA knockdown. (**J**) PGC numbers were significantly reduced following *hh1* or *gli3* knockdown with shRNA. ** represents *p* < 0.01 of two-tailed t-test by comparing with controls (mock-shRNA injection). Scale bar = 50 µm in **A**, **D**, **I**; **B-D** are at the same scale; **F-I’** are at the same scale.

To test functional requirements for Hh signaling in *Nematostella* development, we used shRNA-mediated knockdown and CRISPR/CAS9-directed mutagenesis (Ikmi *et al*., 2014; Kraus *et al*., 2016; He *et al*., 2018). Unfertilized eggs were injected with shRNAs targeting either *hh1* or *gli3* (a transcription factor downstream of Hh signaling) or with two independent *gli3* gRNAs. Using the expression of Vas2 protein and *piwi1* transcript as readouts for PGC identity, we found that PGC specification was significantly inhibited in both knockdown and in presumptive F0 mutant primary polyps (Fig. 5F-J, Fig. 7J-L). These data suggest the Hh signaling pathway is required for *Nematostella* PGC specification.

During Hh signal transduction, binding of Hh ligand to Ptc de-represses the transmembrane protein Smoothened (Smo), which in turn activates a cytoplasmic signaling cascade (Forbes *et al*., 1993; Alcedo *et al*., 1996; Stone *et al*., 1996; van den Heuvel and Ingham, 1996; Bangs and Anderson, 2017). To further test the involvement of Hh signaling in PGC formation, we treated developing animals with the Smo antagonists GDC-0449 or Cyclopamine (McCabe and Leahy, 2015; Sharpe *et al*., 2015). When we treated embryos with either inhibitor, PGC numbers were significantly reduced (Fig. 6). To test Hh requirements for the establishment versus maintenance of PGC identity, we treated developing *Nematostella* with GDC-0449 either during PGC formation (4-8 dpf) or post-PGC formation (8-12 dpf, Fig. S8). PGC formation in 4 to 8 dpf late-planula larvae was significantly inhibited by GDC-0449 treatment (Fig. S8B, compare Ctrl and GDC), while PGC number showed no significant difference when the pathway is inhibited after specification (Fig. S8B, compare Ctrl-Ctrl and Ctrl-GDC). Furthermore, when we compared no-inhibition, continuous-inhibition and released-from-inhibition conditions (Fig. S8B, compare Ctrl-Ctrl, GDC-GDC and GDC-Ctrl), PGC numbers did not vary significantly. These observations suggest that even though initial PGC formation is Hh dependent, the PGC population can be dynamically replenished, potentially through cell proliferation. Additionally, we did not observe PGC migration defects in different combinations of GDC-0449 treatments. At 12 dpf we found that half of treated polyps still showed the expected PGC migration away from the clusters (18 of 29 in Ctrl-Ctrl; 18 of 30 in Ctrl-GDC; 19 of 30 in GDC-GDC; 16 of 30 in GDC-Ctrl). Therefore, Hh signaling is not likely to be involved in PGC migration after their initial specification.

**Fig. 6.**
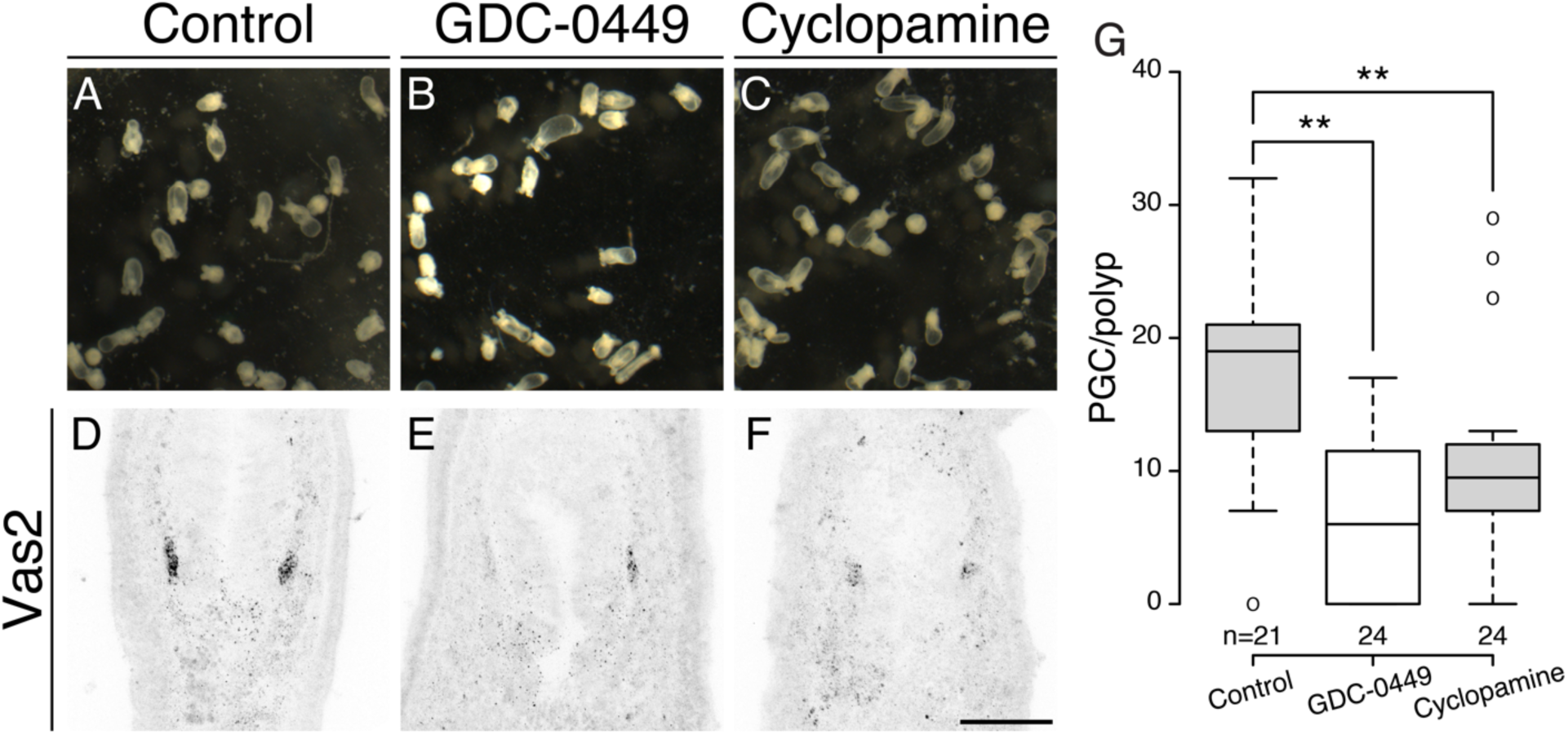
Inhibiting Hh signaling by GDC-0449 or Cyclopamine impairs PGC formation. (**A-C**) The majority of primary polyps do not show visible developmental defects after treatment with GDC-0449 or Cyclopamine from the gastrula stage onward. (**D-G**) Primary polyps treated with GDC-0449 or Cyclopamine formed fewer PGCs than controls. ** represents *p* < 0.01 of two-tailed t-test by comparing with control treatment. Scale bar = 50 µm in **F**. **A-C** are at the same scale; **E-F** are at the same scale.

### Hh signaling regulates endomesodermal patterning and PGC specification

To definitively test the requirements for Hh signaling in *Nematostella*, we used an established CRISPR/Cas9 methodology to mutate *hh1* (Ikmi *et al*., 2014; Kraus *et al*., 2016; He *et al*., 2018). These efforts generated two F1 heterozygous frame-shift lines (a −1 nucleotide frameshift in *hh1*^Δ1^/*+* and a +2 nucleotide frameshift in *hh1*^+2^/*+*; see Materials and Methods). In F2 progeny resulting from crosses between heterozygous F1 siblings, we observed developmental defects in primary polyps wherein body length was reduced by approximately 50% and the four primary tentacles failed to elongate and partially fused together (Fig. 7B-C’). Consistent with a defect in Hh signal transduction, homozygous mutants expressed lower levels of *ptc*, a conserved Hh pathway target gene (Fig. 7D-E’).

**Fig. 7.**
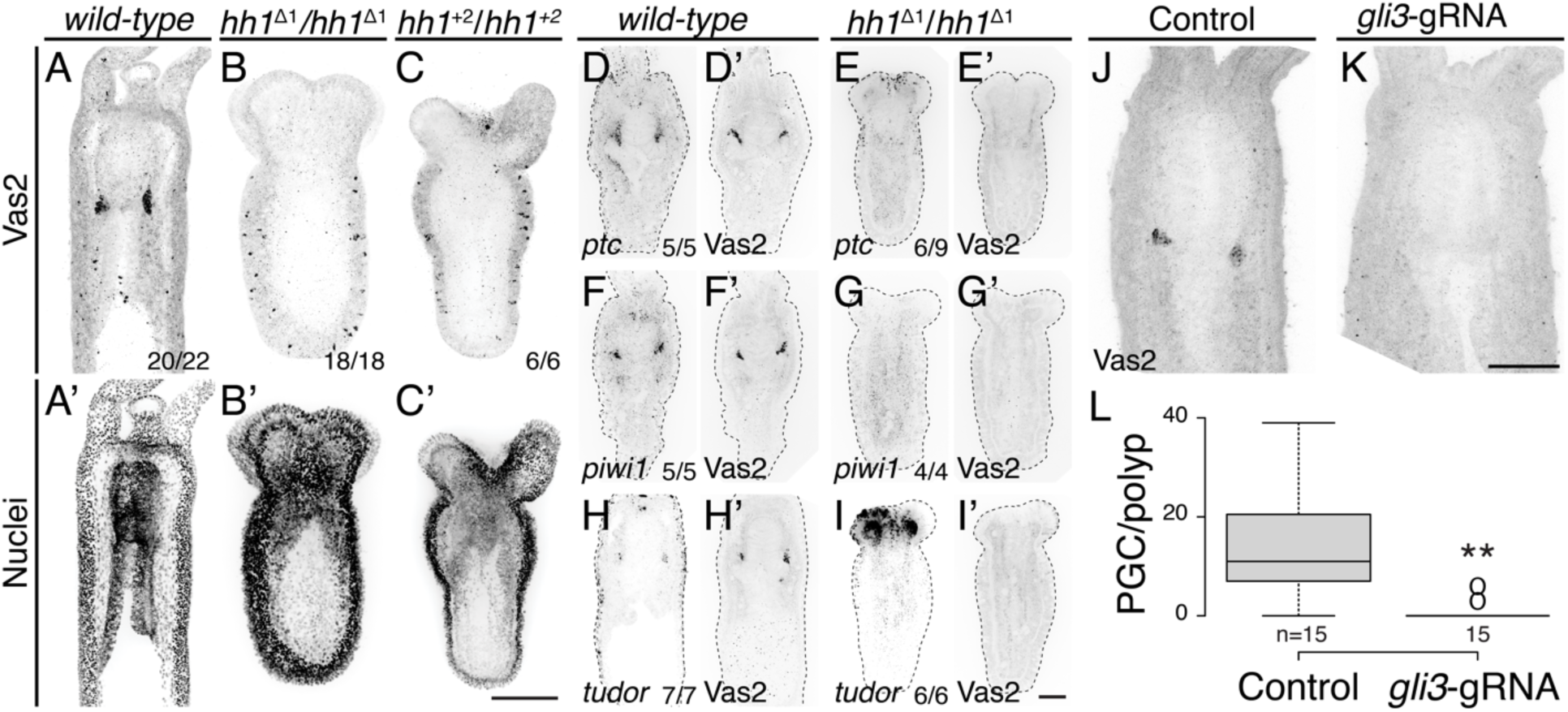
PGC clusters do not form in in *hh1* or *gli3* knockout mutants. (**A-C’**) *hh1* homozygous mutant polyps exhibit a shorter body column and pronounced tentacle defects compared to *wild-type* siblings. Additionally, no Vas2+ PGC clusters were detected in *hh1* homozygous knockout polyps. (**D-E’**) *hh1* mutant polyps show reduced *ptc* expression in the endomesoderm. (**F-I’**) *hh1* mutant polyps lose other PGC markers, including *piwi1* and *tudor*. (**J-L**) Double *gli3* gRNA injected primary polyps showed reduced PGC numbers, as observed in *gli3* shRNA knockdown. **A-C’** are multiple-focal planes projections. **D-I’** are from single-focal plane at the primary mesentery level. Scale bar = 50 µm in **C**, **I’** and **K**; **A-C’** are at the same scale; **E-I’** are at the same scale; **J** and **K** are at the same scale.

Primary polyps homozygous for either *hh1* mutant allele developed primary mesentery-like endomesodermal septa; however, we did not observe Vas2, *piwi1* or *tudor* expressing PGC-like cluster cells (Fig. 7A-I’). Morphological analysis revealed abnormal internal tissue patterning in *hh1* homozygous mutants. The developing pharynx and primary septal filaments are normally connected to body wall ectoderm via endomesodermal tissue (Fig. 8A-A’). In contrast, in *hh1* mutants part of the pharynx and the primary septal filaments were in direct contact with the ectoderm without endomesodermal tissue in between (Fig. 8B-B’). As a result, the eight segments of the larval body plan were abnormally segregated into groups of three and five segments by the pharynx (Fig. 8B-B’). These strong endomesodermal patterning defects were not observed in *hh1* and *gli3* shRNA knockdowns, suggesting PGC formation may require a higher level of Hh signaling activity than endomesoderm patterning. The primary polyp-like *hh1* homozygous mutants passed through gastrulation, suggesting that the pharynx and the endomesoderm likely formed a continuous epithelium (Fig. 7B’ and C’). Consistent with the hypothesis that a pharyngeal signal induces PGC development, *hh1* mutants failed to develop PGCs even though the pharynx associated with the endomesoderm. Nevertheless, without more sophisticated genetic tools, we cannot rule out the possibility that PGC formation was indirectly perturbed by Hh-dependent endomesodermal patterning defects. In either case, we conclude that zygotic signaling activity is required for specification of the putative PGC clusters.

**Fig. 8.**
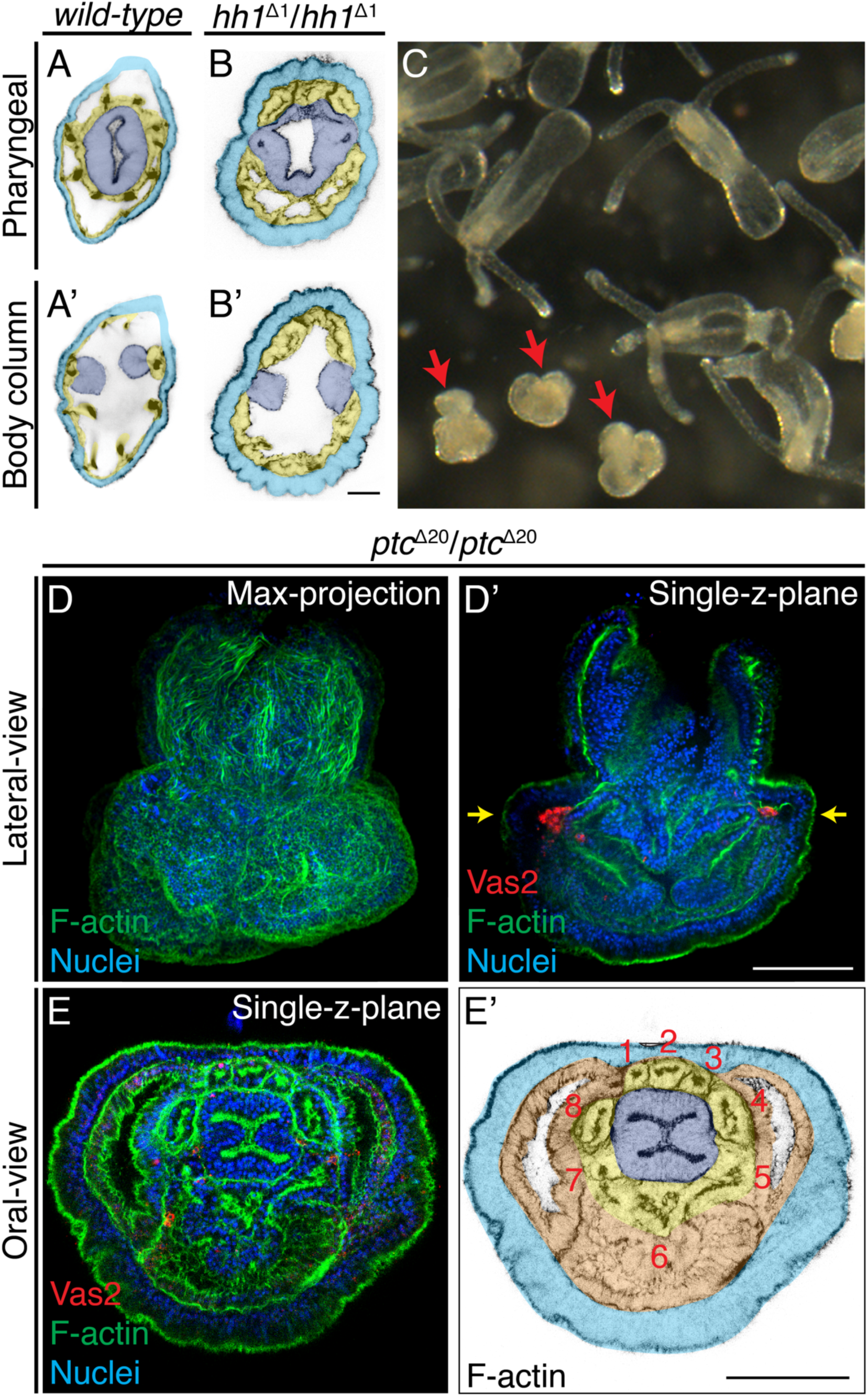
Patterning defects in *hh1* and *ptc* mutants. (**A-B’**) In addition to loss of putative PGCs, *hh1* mutant polyps show endomesodermal patterning defects. Parts of the pharyngeal ectoderm and septal filaments (*navy blue*) abnormally contact the outer epidermis (*azure blue*), without endomesoderm tissue in between (*yellow*). These contacts segregated the normally contiguous eight endomesodermal segments into blocks of three and five segments along the directive axis. (**C**) F2 progeny from a cross between *ptc*^Δ2^/*+* heterozygous siblings. The abnormal mushroom-shaped polyps are indicated by *red* arrows. (**D-D’**) At the primary polyp stage, homozygous *ptc* mutants lack the four primary tentacles and do not develop the normal polyp body plan. (**E-E’**) A single false-colored focal plane taken at the level indicated by yellow arrows in **D’**. Depite significant morphological defects, *ptc* mutants animals develop a pharynx (*navy blue*), eight endomesodermal segments (*yellow*), body wall endomesoderm (*orange*) and putative PGC clusters (labeled by Vas2 immunofluorescence in *red* in **D’**). Scale bar = 20 µm in **B’**; 50 µm in **D’** and **E’**; **A**-**B’** are at the same scale; **D**-**D’** are at the same scale; **E**-**E’** are at the same scale.

### PGC formation in *ptc* mutants may reflect a default Hh activation without the receptor

In bilaterian model systems, Ptc has been shown to serve as a receptor for the Hh ligand and to inhibit the pathway when the ligand is absent (Johnson, Milenkovic and Scott, 2000). To further interrogate the mechanism of PGC specification in *Nematostella*, we generated two *ptc* heterozygous mutant lines (a −2 nucleotide frameshift in *ptc^Δ2^*/+ and a −20 nucleotide frameshift in *ptc^Δ20^*/+, see Materials and Methods). Crosses between *ptc^Δ2^*/+ or *ptc^Δ20^*/+ heterozygous siblings resulted in the expected 25% of homozygous progeny by genotyping, and these developed into abnormal mushroom-shaped polyps which lacked the four primary tentacles (Fig. 8C-D). Detailed morphological examination and Vas2 immunofluorescence revealed that the *ptc* homozygous mutants developed a pharynx, eight endomesoderm mesenteries, and two PGC clusters (Fig. 8D-E’). Combined with the requirement for *hh1*, *gli3* and Smo activity during PGC specification, we propose that the presence of Hh ligand or absence of *ptc* activates the pathway. In turn, zygotic Hh signaling provides permissive conditions for PGC formation at the pharyngeal domain of *Nematostella* endomesoderm.

## Discussion

In this report, we confirm that *Nematostella* putative PGCs form in pharyngeal endomesoderm and provide evidence that these cells delaminate via EMT and migrate through the mesoglea to populate the eight gonad primordia. We also present evidence that putative PGCs form between the expression domains of *hh1* and *ptc* and demonstrate that Hh signaling is required for PGC specification but not PGC maintenance. Because Hh signaling transducers are only expressed zygotically (Matus *et al*., 2008; Lotan *et al*., 2014), these data indicate that *Nematostella* employs an epigenic mechanism to specify PGC fate, which is consistent with the proposed ancestral mechanism for metazoan PGC specification (Extavour and Akam, 2003). It remains possible that maternally inherited germline determinants still play some essential role in PGC specification and that zygotic Hh activity serves to augment their function. In this combined maternal-zygotic scenario, the mechanism of *Nematostella* PGC formation would not neatly fit within either preformation or epigenesis and instead fall within the continuum between either extreme, similar to sea urchin PGCs (Seervai and Wessel, 2013).

### Hh pathway activity in *ptc* mutants

In many bilaterian model organisms *ptc* is a transcriptional target of Hh signaling and serves as both a receptor and negative regulator of pathway activity (Briscoe and Thérond, 2013; Bangs and Anderson, 2017). We sought to functionally dissect Hh signaling in *Nematostella* and leveraged CRISPR/Cas9 mutagenesis to generate both *hh1* and *ptc* mutants. While *hh1* mutants lacked putative PGC cell clusters (Fig. 7), to our surprise these cells formed properly in *ptc* mutant animals (Fig. 8). This finding could be consistent with three possible scenarios: **1**) The existence of residual receptor activity due to allele-specific effects or potential redundancy with an unannotated orthologue elsewhere in the genome; **2**) An indirect role for Hh in PGC specification; **3**) A default repressive role for Ptc in the specification of pre-patterned PGC clusters. Because inhibiting Hh activity by disrupting either Smo or *gli3* also disrupted PGC formation, we suggest that *ptc* most likely serves as a default inhibitor of Hh activity in *Nematostella*. Based on our combined data, we propose that the pharyngeal ectoderm releases Hh ligand to inhibit Ptc-dependent repression of PGC fate in neighboring endomesoderm. This reasoning would suggest that the PGC clusters are pre-patterned by other yet-identified extracellular signals, and that the role of Hh activity may be to provide a spatial or temporal cue to trigger their maturation.

### Direct versus indirect roles for Hh activity in PGC specification

To our knowledge, Hh signaling has not been directly implicated in PGC specification in previous studies of established bilaterian systems. Nevertheless, a requirement for Hh signaling during *Nematostella* PGC formation is supported by three lines of evidence: **1**) *hh1* and *gli3* shRNA knockdowns (Fig. 5); **2**) Smo inhibition assays (Fig. 6); and **3**) *hh1* and *gli3* CRISPR/Cas9 mutagenesis (Fig. 7). In developing primary mesenteries, PGCs are specified in endomesoderm cells that lie in close proximity to the *hh1* expression domain in adjacent pharyngeal ectoderm (Fig. 5). Even in the absence of primary mesenteries in *Anthox6a* mutants and *Gbx* knockdown juveniles (Fig. S7; He *et al*., 2018), PGCs still develop from endomesodermal cells in proximity to *hh1*-expressing ectodermal septal filament cells. Interestingly, while *hh1* expression seems to be restricted to the pharyngeal ectoderm and septal filaments, we observed broad endomesodermal patterning defects in *hh1* mutants (Fig. 8). This phenotype was not seen in either knockdown experiments or inhibitor assays where PGC specification was nevertheless inhibited (Fig. 5 and Fig. 6). This could suggest that the PGC defects in *hh1* mutants are not an indirect result of aberrant endomesodermal patterning. Still, looking forward, genetic tools that allow the discrimination between cell autonomous and cell non-autonomous genetic requirements will be required to definitively rule out whether PGC formation is directly or indirectly regulated by Hh signaling.

### Future perspectives

In this report, we provide an initial framework demonstrating an epigenic PGC formation mechanism in *Nematostella vectensis*, a representative early-branching animal. Because Hh signaling components are found in choanoflagellates and the Hh pathway dictates neighbor cell fates throughout developing metazoans (Adamska *et al*., 2007; King *et al*., 2008), it remains possible that this pathway distinguishes germline and soma in other animals as well. Alternatively, the requirement for Hh signaling in *Nematostella* PGC formation could be a lineage-specific feature of sea anemones, which could be addressed through a broad sampling of PGC development in diverse anthozoan cnidarians.

## Materials and methods

### Animal husbandry

Maintenance and spawning of *N. vectensis* followed previously established protocols (Stefanik, Friedman and Finnerty, 2013). Embryos were cultured at 24 °C for consistent developmental staging.

### Generation of anti-Vas2 antibody

A His-tagged antigen for raising polyclonal Rabbit-anti-Vas2 antibody was designed, synthesized and purified by GenScript (Piscataway, NJ). The antigen sequence (MCFKCQQTGHFARECPNESAAGENGDRPKPVTYVPPTPTEDEEEMFRSTIQQGINFEKYDQIEVLVSGNNPVRHINSFEEANLYEAFLNNVRKAQYKKPTP<u>HHHHHH</u>) partially encompasses the last zinc finger domain and the DEAD-like helicase domain of Vas2. In brief, the rabbit was immunized 3 times before checking the antibody titer, boosted once more and sacrificed for the whole serum. The serum was affinity purified, and the polyclonal antibody stock concentration is 0.4 mg/mL.

### Whole-mount immunofluorescence

Our immunohistochemical staining protocol generally followed Genikhovich and Technau (2009), with the following modifications: samples were blocked in 5% goat serum diluted in PBS with 0.2% Triton X-100 (PTx) and 10% DMSO for increasing antibody penetration. Samples were then incubated in 1:1000 diluted stock of Rabbit-anti-Vas2 antibody in PTx with 0.1% DMSO and 5% goat serum at 4 °C overnight. After six washes with PTx for at least 20 minutes each at room temperature, samples were incubated with Alexa Fluor Goat-anti-Rabbit secondary antibodies (Thermo Fisher Scientific; Waltham, MA) at 1:500 dilution in PTx with 5% goat serum at 4 °C overnight. If desired, during secondary antibody incubation, samples were counter-stained for F-Actin with Phalloidin at 1:200 dilution (Thermo Fisher Scientific) and nuclei with either 1 µg/mL of Hoechst-34580 (Sigma-Aldrich; St. Louis, MO; Cat. No. 63493) or 1:5,000 diluted SYBR™ Green I (Thermo Fisher Scientific; Cat. No. S7567). After several washes, samples were serially immersed in a final solution of 80% of glycerol in PTx. Alternatively, we dehydrated samples in isopropanol and immersed in BABB (one part of benzyl alcohol and two parts of benzyl benzoate) to clear the tissue, allowing up to 400 µm imaging depth (Wan *et al*., 2018).

### Whole-mount fluorescent *in situ* hybridization (FISH)

To clone target genes, purified total RNA was reverse transcribed into cDNA by ImProm-II™ Reverse Transcription System (Promega; Madison, WI; Cat. No. A3800). Target gene fragments were first amplified from a mixed cDNA library of planula larva and primary polyps. Primers are listed in Table S1. We adopted a ligation-independent pPR-T4P cloning method (Newmark et al., 2003) to generate plasmids with probe templates and confirmed the positive clones by sequencing. We then PCR amplified the DNA template fragments using the AA18 (CCACCGGTTCCATGGCTAGC) and PR244 (GGCCCCAAGGGGTTATGTGG) primers, which flank the T7 promoter and the target gene sequence. After purifying DNA templates, we synthesized DIG-labeled RNA probes with the DIG RNA labeling mix (Sigma-Aldrich; Cat. No. 11277073910) and T7 RNA polymerase (Promega; Cat. No. P2077). Sample preparation, probe hybridization and signal detection followed established protocols (Steinmetz *et al*., 2017; He *et al*., 2018). The probe working concentration was 0.5 ng/µL for all genes.

For double FISH, we synthesized fluorescein-labeled RNA probes with the Fluorescein RNA labeling mix (Sigma-Aldrich; Cat. No. 11685619910) and hybridized together with a DIG-labeled probe of another gene. After detecting the first probe signal with TSA® fluorescein reagent (PerkinElmer; Waltham, MA) and several washes with TNT buffer, we quenched peroxidase activity by incubating samples in 200 mM NaN_3_/TNT for 1 hour. Samples were then washed six times with TNT for at least 20 min each and then subjected to second-round probe detection by either anti-DIG-POD Fab fragments (Sigma-Aldrich; Cat. No. 11207733910) or Anti-Fluorescein-POD Fab fragments (Sigma-Aldrich; Cat. No. 11426346910).

### Short hairpin RNA knockdown

shRNA design, synthesis and delivery followed the protocol of He et al. (2018) with the following modification: A reverse DNA primer containing the shRNA stem and linker sequence was annealed with a 20nt T7 promoter primer (TAATACGACTCACTATAGGG). The annealed, partially double stranded DNA directly served as template for *in vitro* transcription. We tested knockdown efficiency with shRNA produced by this modified method by targeting *β-catenin* and *dpp* shRNA, and found the same phenotypic penetrance as the previous method (He *et al*., 2018; Karabulut *et al*., 2019). To control for shRNA toxicity, we injected 1000 ng/µL eGFP shRNA and did not observe noticeable developmental defects. All shRNA working solutions were prepared at 1000 ng/µL and the sequences are listed in Table S2. By 8 dpf, primary polyps were fixed to assay PGC development.

### Generation of mutant lines by CRISPR/Cas9 mutagenesis

*hh1* and *ptc* mutant lines were generated using established methods (He *et al*., 2018). In brief, to generate F0 founders, we co-injected 500ng/µl of gRNA (sequences listed in Table S3) and 500ng/µl of SpCas9 protein into unfertilized eggs. Mosaic F0 founders were then crossed with wild-type sperm or eggs to create a heterozygous F1 population. When the F1 polyps reached juvenile stage, we genotyped individual polyps by cutting tentacle samples; resultant alleles are as described in Table S4. Heterozygous carriers of insertion/deletion-induced frame-shift alleles were crossed to generate homozygous mutants. The phenotypes and genotypes of this F2 population followed mendelian inheritance and were subjected to further analysis. In progeny resulting from a *hh1^Δ1^/+* cross, the observed phenotypic ratio of wild-type and mutant primary polyps was 948:343, close to the expected mendelian ratio. Progeny from heterozygous crosses were also randomly genotyped and confirmed to follow the expected 1:2:1 ratio (*+*/*+*: *hh1*^Δ1^/*+*: *hh1*^Δ1^/*hh1*^Δ1^ = 6:14:8 and *+*/*+*: *hh1*^+2^/*+*: *hh1^+^*^2^/*hh1*^+2^ = 7:16:6). A similar strategy was used to analyze *ptc* mutants. *ptc* mutant genotypes also followed mendelian segregation (*+*/*+*: *ptc*^Δ2^/*+*: *ptc*^Δ2^/*ptc*^Δ2^ = 5:17:8; (*+*/*+*: *ptc*^Δ20^/*+*: *ptc*^Δ20^/*ptc*^Δ20^ = 7:12:11). These results suggest the phenotypes observed in the *hh1* and *ptc* mutant lines result from single locus mutations.

### Inhibitor treatments

GDC-0449 (Cayman Chemical; Ann Arbor, MI; Cat. No. 13613) and Cyclopamine (Cayman Chemical; Cat. No. 11321) were diluted in DMSO as 50 mM and 10 mM stocks, respectively. These stocks were diluted 1:2000 in 12 ppt filtered artificial sea water (FSW) to generate working solution: 25 µM GDC-0449 and 5 µM Cyclopamine. 1:2000 diluted 100% DMSO (final 0.05%) was applied as a control. All treatments were protected from light, incubated at 24 °C, and replaced with fresh working solutions every day.

### Imaging and quantification

For confocal imaging, we used a Leica TCS Sp5 Confocal Laser Scanning Microscope or a Nikon 3PO Spinning Disk Confocal System. Bright field images were acquired using a Leica MZ 16 F stereoscope equipped with QICAM Fast 1394 camera (Qimaging; Surrey, BC, Canada). The brightness and contrast of images were adjusted by Fiji, and the PGC number of individual polyp was manually quantified by the Cell Counter Macro or automatically by blurring and masking the Vasa2 signal to find the cluster and 3D peak finding of the DAPI nuclei within the cluster (http://github.com/jouyun/smc-macros). Serial z section images of *N. vectensis* pharyngeal structures were reconstructed as a 3D movie (Mov. S1) by Imaris 8.3 (Bitplane, Concord, MA). Figures of this report were generated using Adobe Illustrator 2019.

## Acknowledgements

The authors would like to thank Gibson lab members for their suggestions and critical readings of the manuscript. Additionally, we thank the Stowers Institute Core Technology Centers, in particular Aquatics for *Nematostella* husbandry, Molecular Biology for sequencing and genotyping, and Histology for help with histological sectioning. This work was funded in its entirety by the Stowers Institute for Medical Research.

---

**Table S1.**
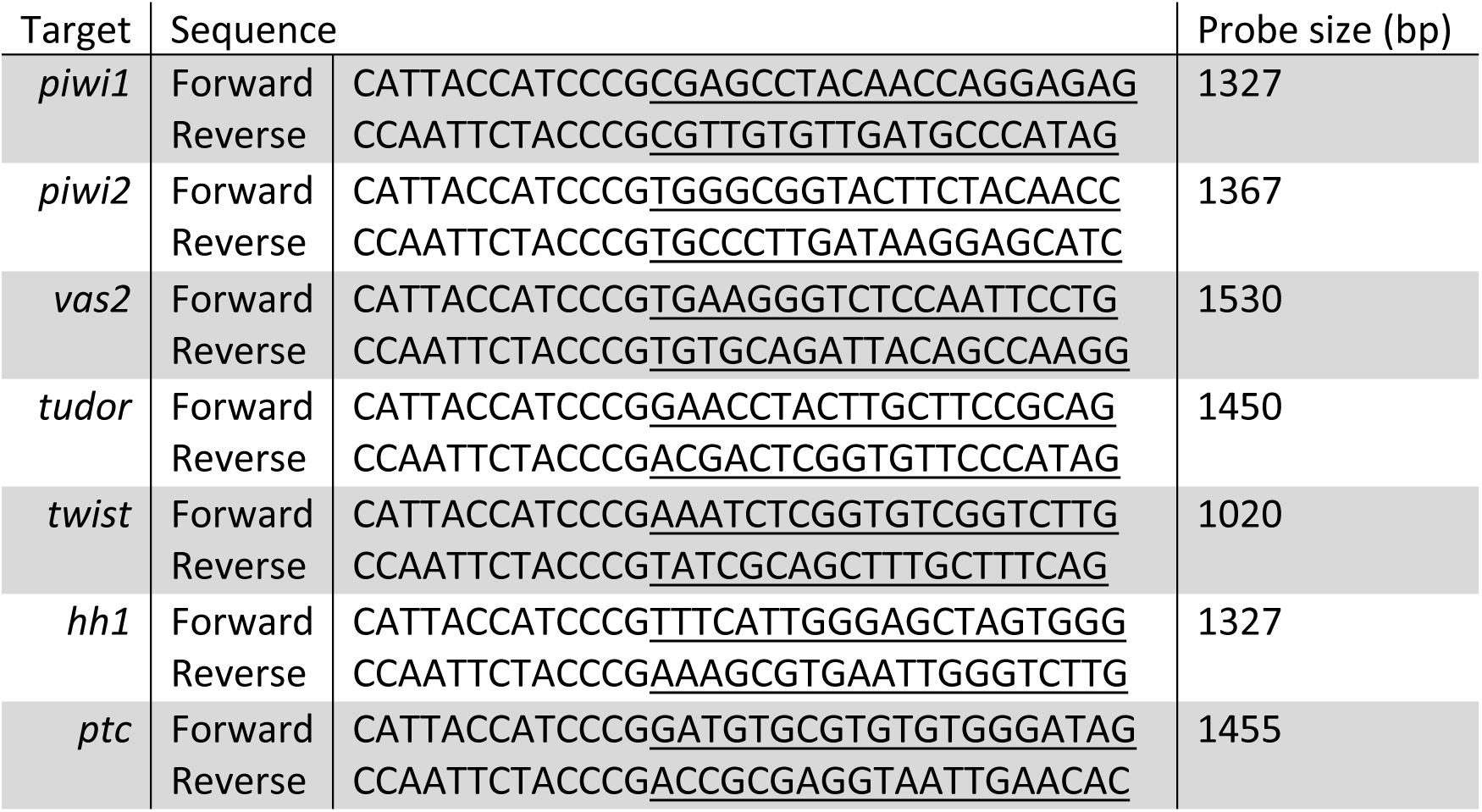
Primer sets for cloning probe templates into pPR-T4P (gene-specific regions are underlined)

**Table S2.**
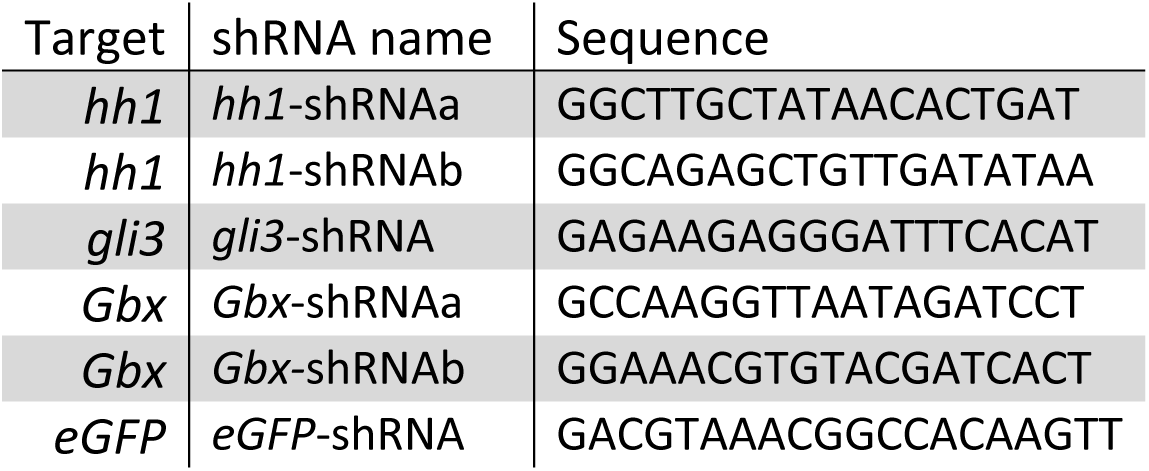
shRNA sequences.

**Table S3.**
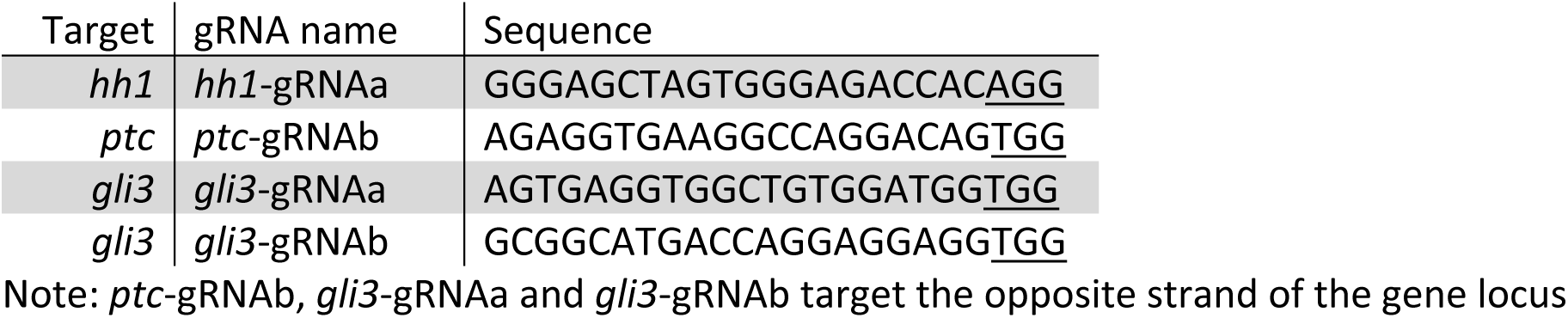
gRNA sequence for CRISPR/Cas9 mutagenesis (PAM domains are underlined)

**Table S4.**
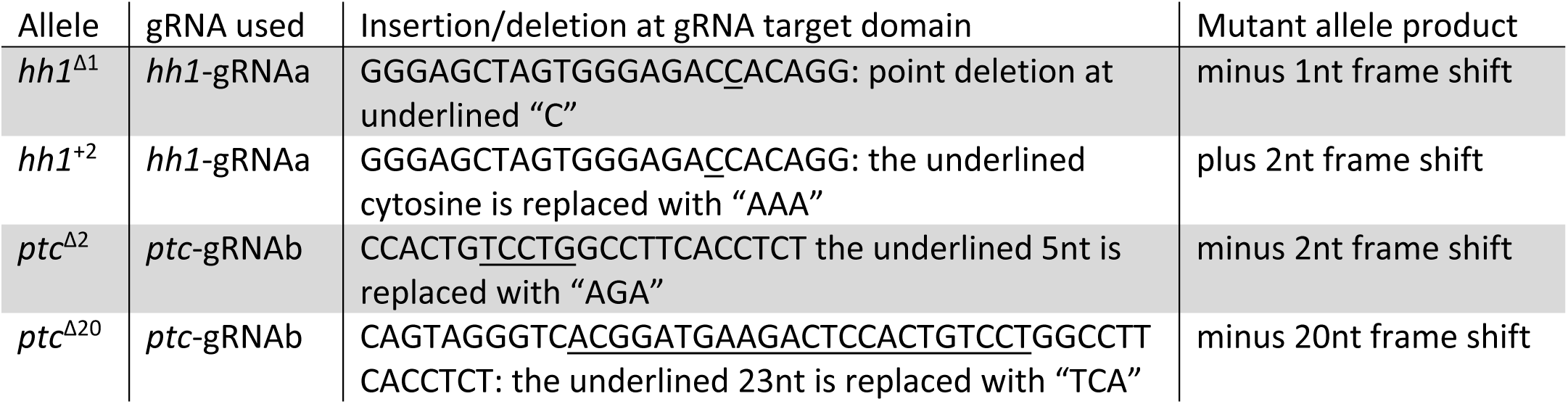
Mutant allele descriptions.

**Fig. S1.**
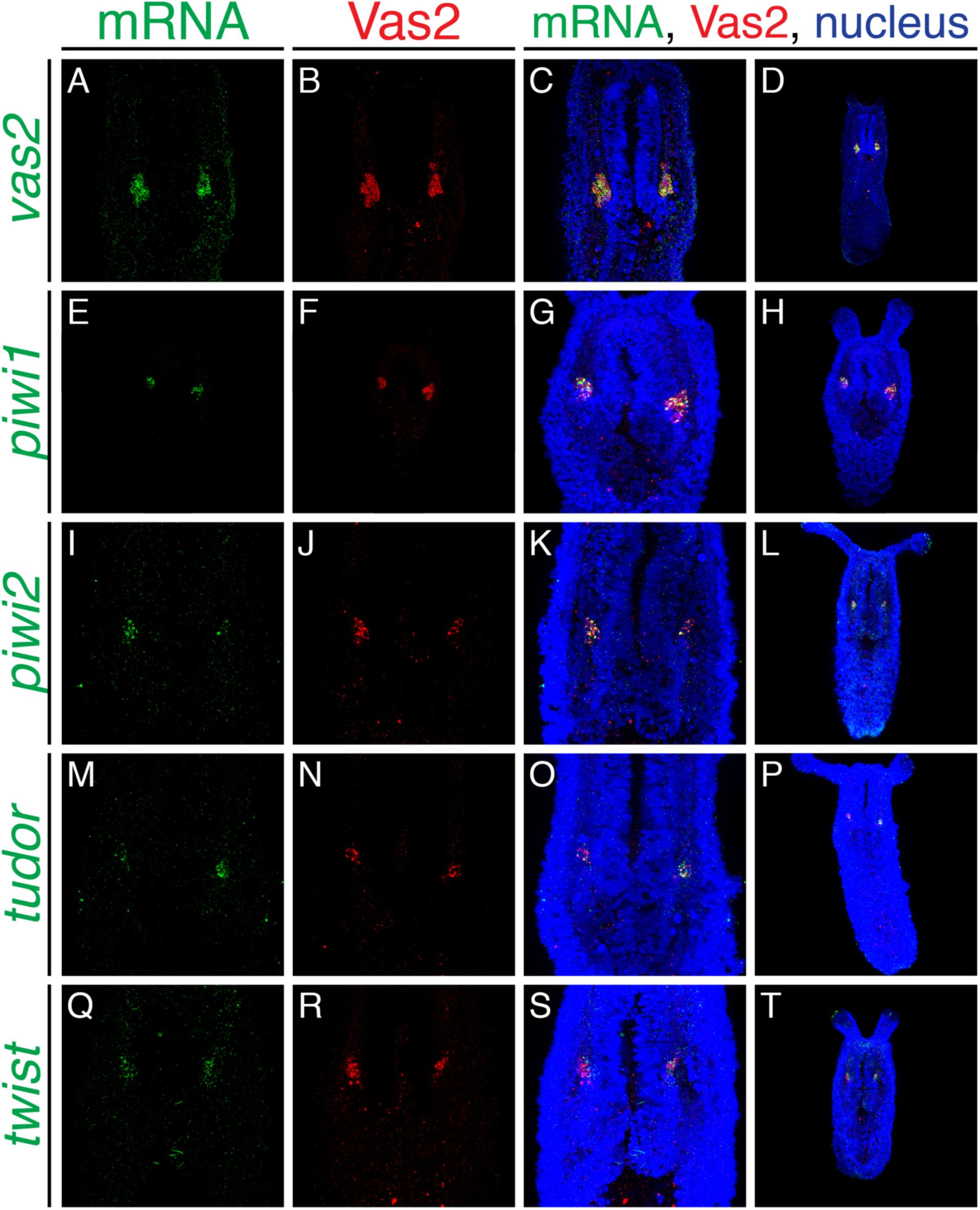
Conserved germline and EMT-related genes are expressed in putative PGC clusters. Fluorescent *in situ* hybridization (FISH, *green*) of *vas2* (**A**), *piwi1* (**E**), *piwi2* (**I**), *tudor* (**M**) and *twist* (**Q**) show enhanced expression within PGC cell clusters (Vas2-immunofluorescence, *red*).

**Fig. S2.**
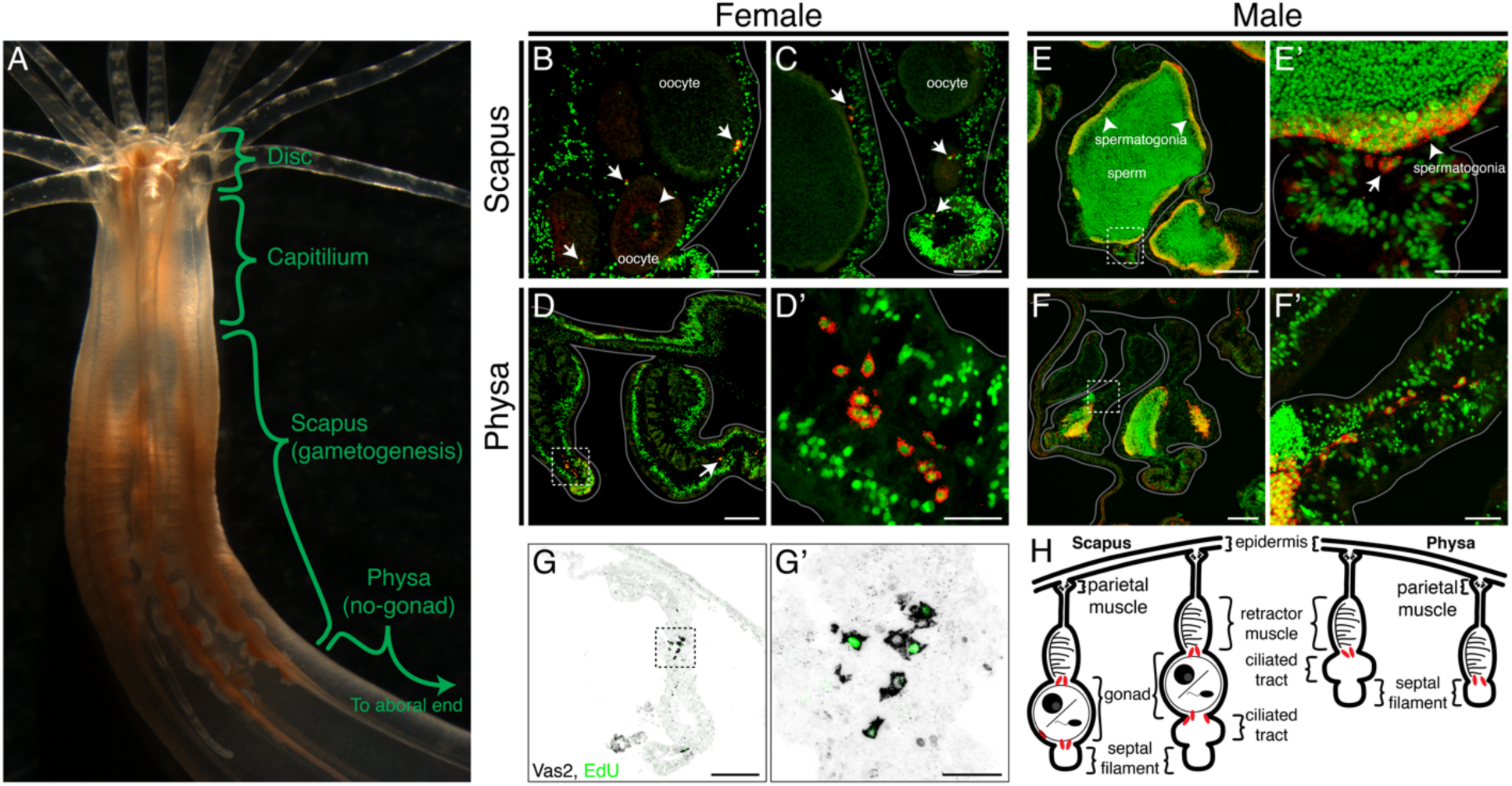
Adult PGC-like lineages localize adjacent to the mature gonad and migrate aborally. (**A**) Along the oral-aboral axis, adult *Nematostella* can be regionally subdivided into the disc (mouth and tentacle base), capitillium (pharynx), scapus (gametogenic region) and physa (non-gametogenic region; Williams, 1979). (**B-F**’) Female and male cross sections were immunolabeled with Vas2 (*red*) and counter-stained for nuclei (*green*). PGC-like cells (*arrows* in **B-C** and **E’**) localize next to the maturing oocytes and sperm stem cells (spermatogonia, *arrowheads* in **E-E’**), which divide and give rise to sperm in the sperm cyst lumen. Note there are Vas2 puncta surrounding the nuclei of maturing oocytes (*arrowhead* in **B**) and spermatogonia. (**D-D’**, **F-F’ and G-G’**) PGC-like cells also localize aborally in the physa, the site of occasional asexual fission (Hand and Uhlinger, 1995; Burton and Finnerty, 2009). Aboral migration may thus ensure fertility in asexually-produced progeny. (**G-G’**) PGC-like cells incorporate EdU (*green*), consistent with proliferative activity. (**H**) Schematic diagrams depict the cross-sectional localization of PGC-like cells (*red*) in adult mesenteries. Scale bar = 50 µm in **B** and **C**; 100 µm in **D**, **E**, **F** and **G**; 20 µm in **D’**, **E’**, **F’**, and **G’**.

**Fig. S3.**
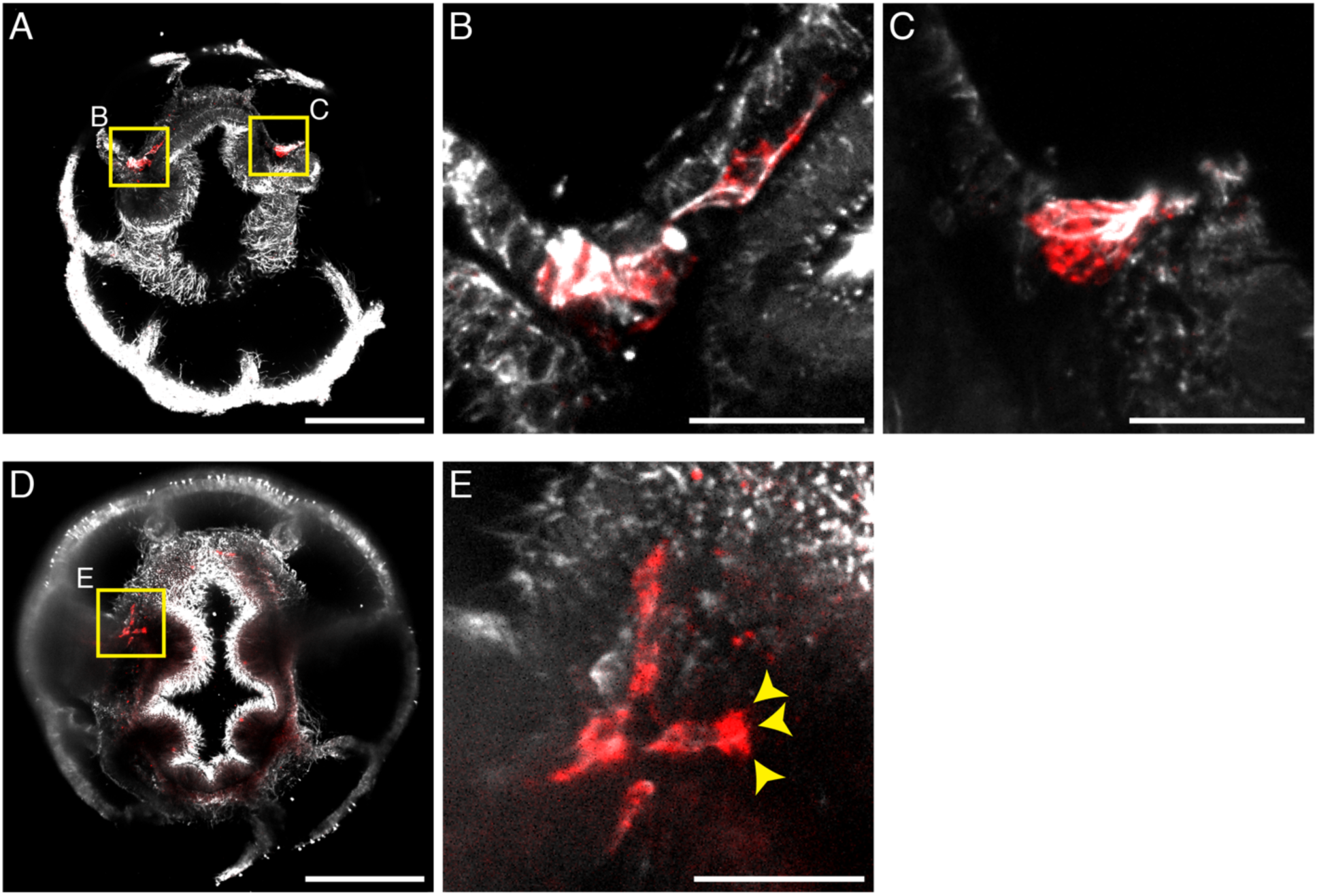
Migratory PGCs show fibroblast morphology and filopodia. (**A**-**E**) Representative images show the microtubule cytoskeleton (α-Tubulin, *grey*) of PGCs (*red*) and their filopodia-like protrusions (*yellow arrowheads*, **E**). Scale bar = 100 µm in **A** and **D**; 20 µm in **B**-**C** and **E**.

**Fig. S4.**
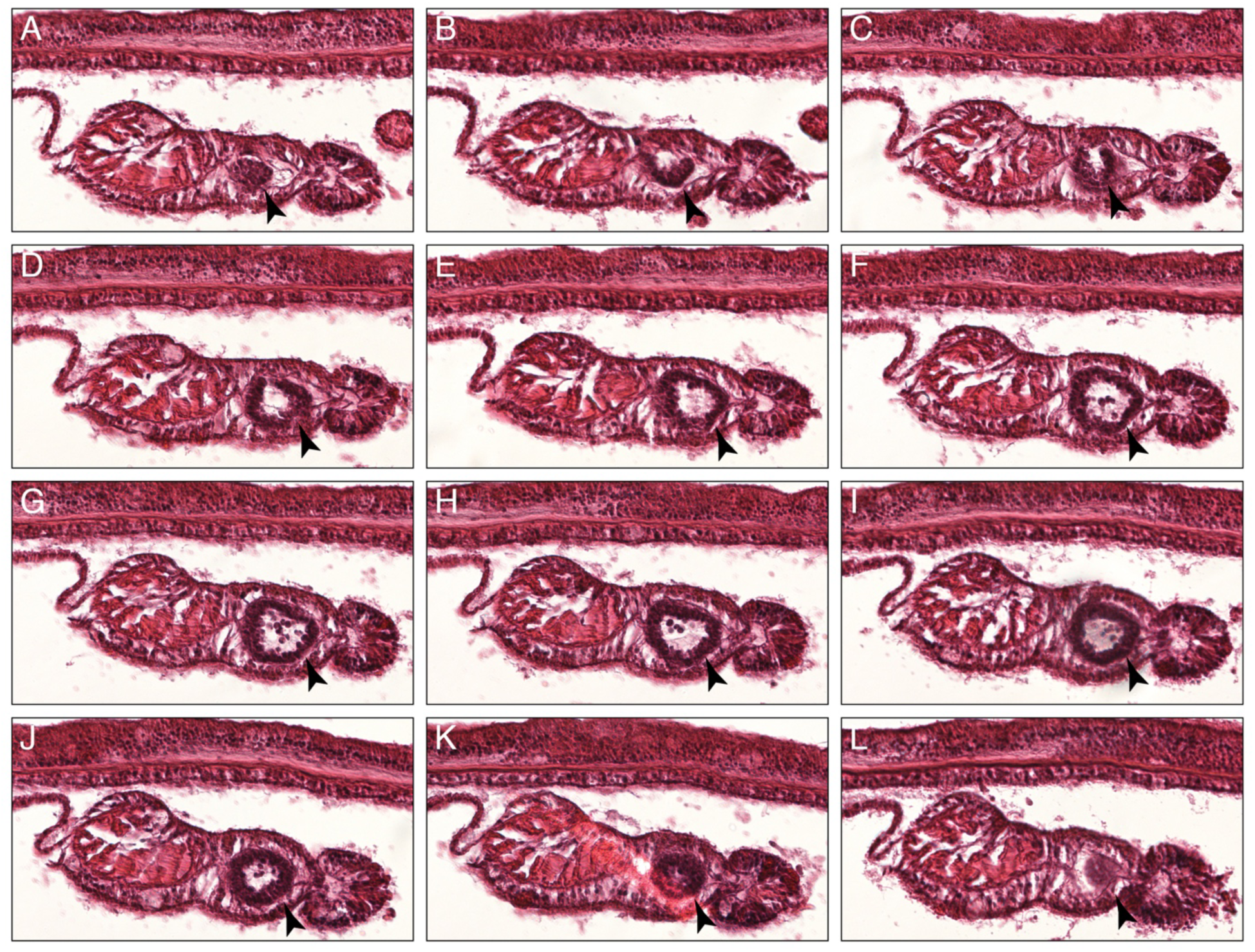
Serial cross sections of a juvenile male mesentery. Germline stem cells (spermatogonia) form cysts (*black arrowheads*) and give rise to maturing sperm in the cyst lumen.

**Fig. S5.**
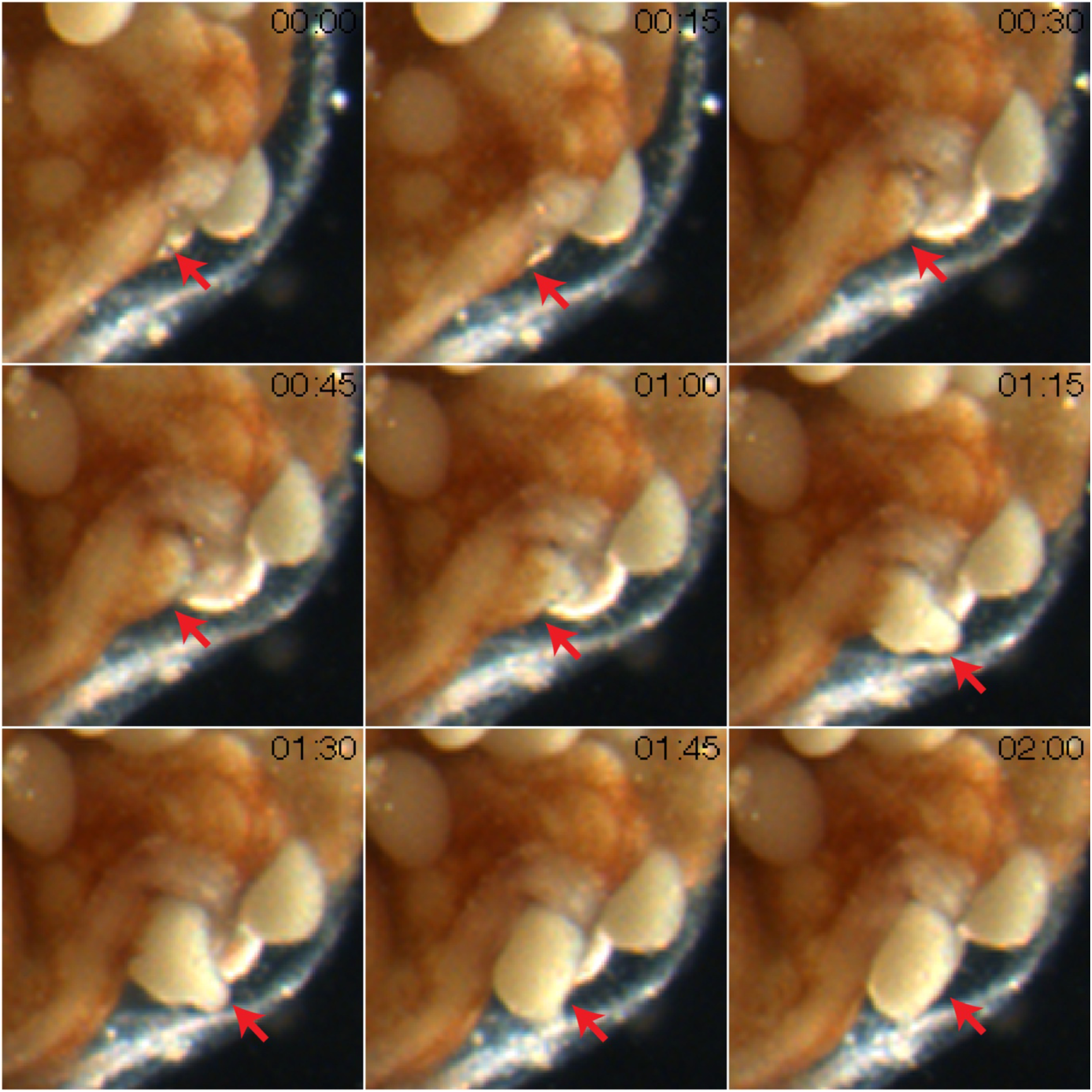
Serial time frames of an oocyte rupturing out of gonad epithelium during spawning. The frames correspond to the green dot-labeled oocyte of Supplementary MOV. 2, 13:00 to 15:00.

**Fig. S6.**
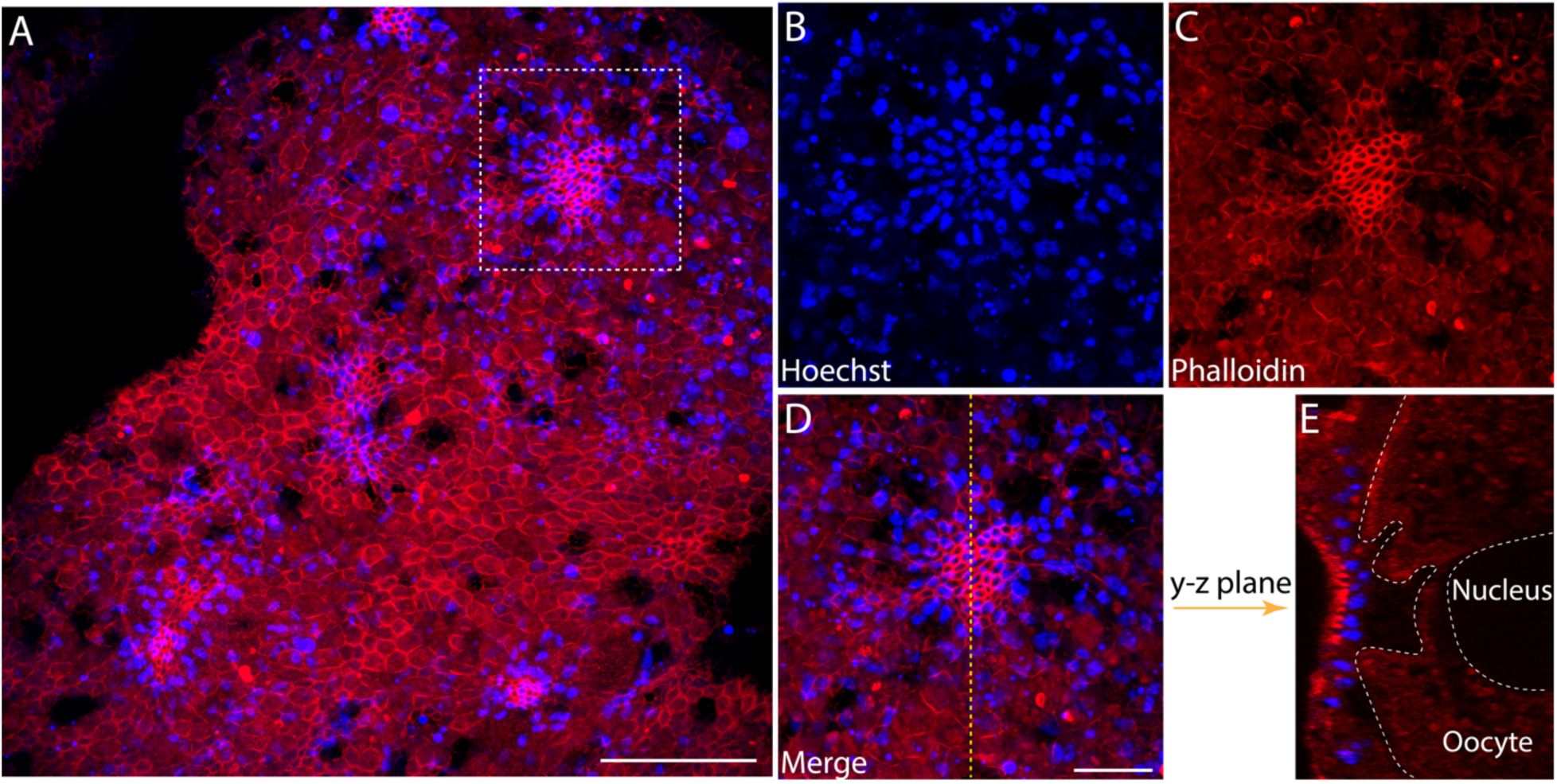
Trophocytes aggregate on the female gonad epidermis and concentrate above the maturing oocyte. (**A**) Apical view of female gonad epidermis. (**B-D**) Single channel images of the box in **A**. Specialized gonad epithelial cells, trophocytes, form trophonema and enrich F-actin on the apical pole, as labelled by Phalloidin in **C**. (**E**) Optical cross section taken along the dotted line in **D**, showing that the trophonema directly overlays a developing oocyte in the gonad lumen. Scale bar = 50 µm in **A**; 20 µm in **D**. **B-E** are at the same scale.

**Fig. S7.**
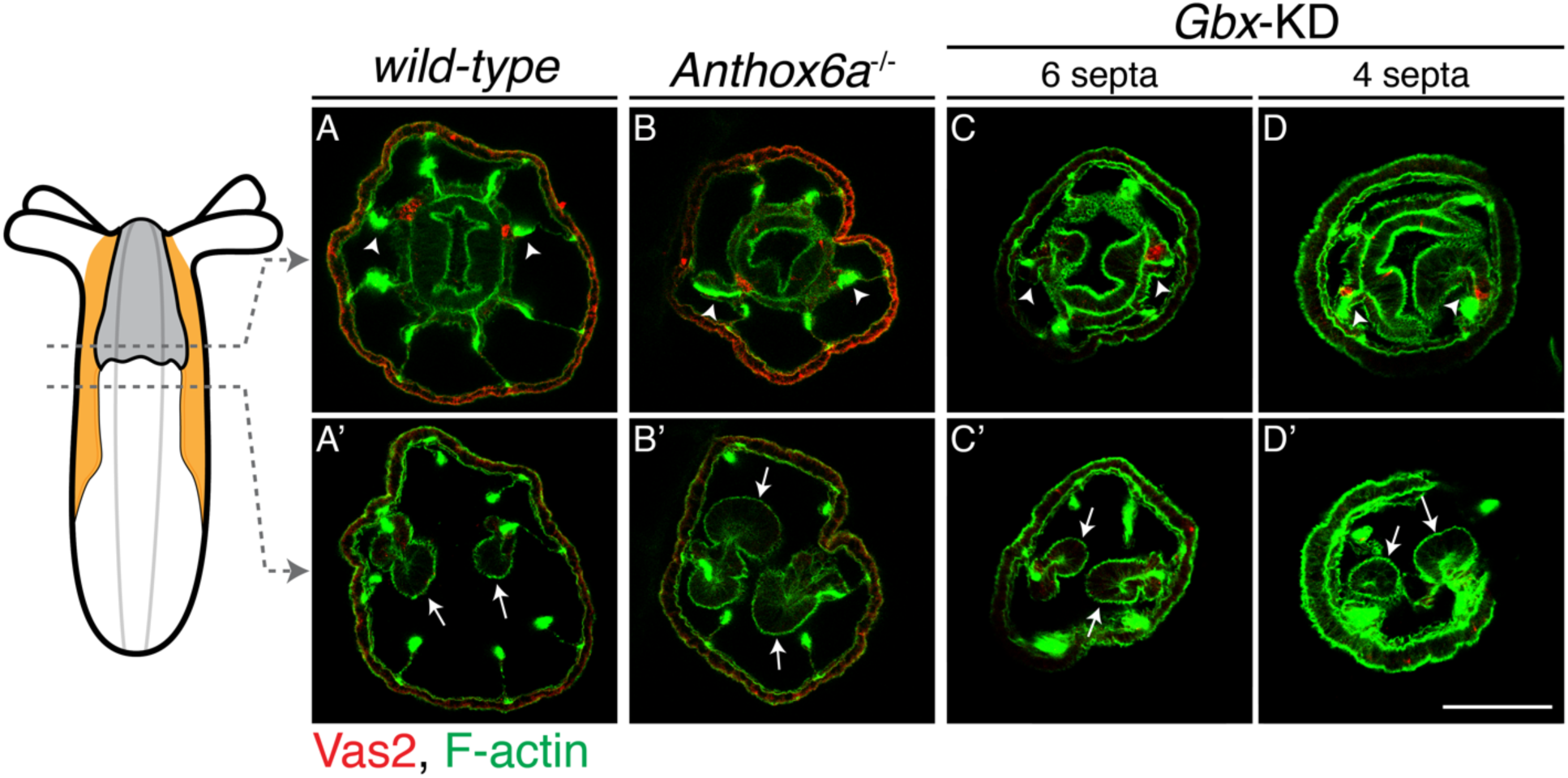
PGC clusters are specified on mesenteries with primary septal filaments of wild-type, *Anthox6a* mutant and *Gbx* shRNA knockdown primary polyps. Although primary mesenteries are missing in *Anthox6a* mutants or *Gbx* shRNA knockdown primary polyps, PGCs (*red*) are specified on the mesenteries (*arrowheads*) where primary septal filaments attach (*arrows*). Scale bar = 50 µm in **D’**. All images are at the same scale.

**Fig. S8.**
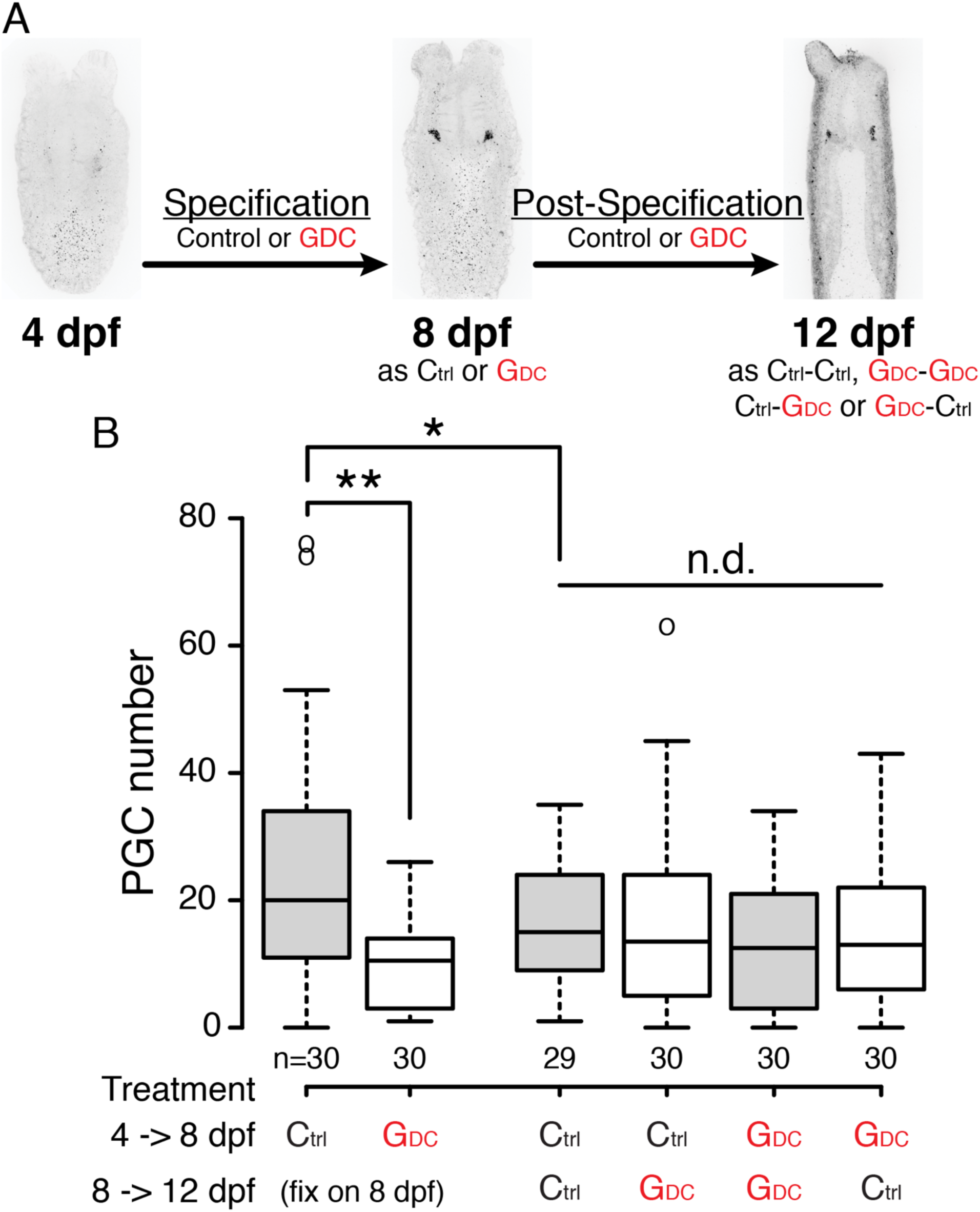
The Hh signaling pathway does not affect PGC behaviors after formation. (**A**) Design of Hh signaling inhibitor GDC-0449 treatments: 4 to 8 dpf larvae were tested for PGC specification, and 8 to 12 dpf polyps were tested for PGC behaviors post-specification. Ctrl: DMSO treatment. GDC: 25 µM GDC-0449 treatment. (**B**) Quantification of PGC numbers after treatments. During PGC specification (4-8 dpf), GDC-0449 resulted in significantly less PGC formation than control. After PGCs are specified (after 8 dpf), PGC number does not show any statistical difference among treatments. ** represents *p* < 0.01 of two-tailed t-test by comparing with DMSO control. n.d.: no-statistically significant difference.

**Sup. Mov.1. 3D reconstruction of pharyngeal structures.** PGCs (*magenta*) are specified on the epithelium of the two primary mesenteries, close to the pharynx (*cyan* cells in the center). The endomesodermal nuclei are pseudocolored in *blue*, and the ectodermal nuclei are in *cyan*.

**Sup. Mov.2. Oocyte spawning time lapse movie.** Nine representative oocytes were tracked by colored dots when rupturing out of the mesentery epithelium. In normal spawning, oocytes mix with egg jelly inside the female body column before the egg mass is pushed out of the pharynx and fertilized *ex vivo*. The time lapse video was taken at 15 sec/frame. The movie is at 30X speed.

